# The *Setaria viridis* genome and diversity panel enables discovery of a novel domestication gene

**DOI:** 10.1101/744557

**Authors:** Sujan Mamidi, Adam Healey, Pu Huang, Jane Grimwood, Jerry Jenkins, Kerrie Barry, Avinash Sreedasyam, Shengqiang Shu, John T. Lovell, Maximilian Feldman, Jinxia Wu, Yunqing Yu, Cindy Chen, Jenifer Johnson, Hitoshi Sakakibara, Takatoshi Kiba, Tetsuya Sakurai, Rachel Tavares, Dmitri A. Nusinow, Ivan Baxter, Jeremy Schmutz, Thomas P. Brutnell, Elizabeth A. Kellogg

**Affiliations:** HudsonAlpha Institute for Biotechnology, Huntsville, Alabama, USA; Donald Danforth Plant Science Center, 975 North Warson Road, St. Louis, MO 63132, USA; Department of Energy Joint Genome Institute, Lawrence Berkeley National Laboratory, 1 Cyclotron Road, Berkeley, CA 94720, USA; RIKEN Center for Sustainable Resource Science, Tsurumi, Yokohama 230-0045, Japan; BASF Corporation, 26 Davis Dr., Durham, NC 27709, USA; USDA-ARS Temperate Tree Fruit and Vegetable Research Unit, 24106 N. Bunn Rd., Prosser, WA 99350, USA; Biotechnology Research Institute, Chinese Academy of Agricultural Sciences, Beijing 100081, China; Graduate School of Bioagricultural Sciences, Nagoya University, Nagoya 464-8601, Japan; Multidisciplinary Science Cluster, Kochi University, Nankoku, Kochi, 783-8502, Japan; Biology Department, University of Massachusetts, Amherst, MA 01003 USA

**Author notes:** These authors contributed equally to this paper (Sujan Mamidi, Adam Healey, Pu Huang).

## Abstract

Diverse wild and weedy crop relatives hold genetic variants underlying key evolutionary innovations of crops under domestication. Here, we provide genome resources and probe the genetic basis of domestication traits in green millet (*Setaria viridis*), a close wild relative of foxtail millet (*S. italica*). Specifically, we develop and exploit a platinum-quality genome assembly and *de novo* assemblies for 598 wild accessions to identify loci underlying a) response to climate, b) a key ‘loss of shattering’ trait that permits mechanical harvest, and c) leaf angle, a major predictor of yield in many grass crops. With CRISPR-Cas9 genome editing, we validated *Less Shattering1* (*SvLES1*) as a novel gene for seed shattering, which is rendered non-functional via a retrotransposon insertion in *SiLes1*, the domesticated loss-of-shattering allele of *S. italica*. Together these results and resources project *S. viridis* as a key model species for complex trait dissection and biotechnological improvement of panicoid crops.

## Introduction

Warm season C_4_ grasses (subfamily Panicoideae) such as maize, sorghum and most species of millet are mainstays of industrial and small-holder agriculture, and include most major biofuel feedstocks. Crucially, C_4_ photosynthesis is most productive under the hot and dry conditions that are predicted to become more prevalent over the coming decades^1–3^. While contemporary breeding of Panicoideae crops targets a trade-off between stress tolerance and yield, de novo domestication of wild Panicoideae species may provide an alternative path to develop novel bioproducts, fuels and food sources. However, the plants in the Panicoideae are known for long lifespans and large or complex genomes, creating the need for a practical experimental model that can be used for rapid discovery of gene structure and function and biotechnological improvement of related crops.

*Setaria viridis* (green foxtail) has emerged as such a model^4–6^. Plants are small (Figure 1a), diploid, have a short life cycle (seed to seed in 8-10 weeks), a small genome (ca. 500 Mb), and are self-compatible, with a single inflorescence that often produces hundreds of seeds. Transformation is efficient, and amenable to CRISPR-Cas9 mediated mutagenesis. These features have enabled use of *S. viridis* for studies of photosynthetic mechanisms^7–11^, drought tolerance^12^, cell wall composition^13–15^, floral and inflorescence development^16, 17^, leaf anatomy^18^, secondary metabolism^19^, plant microbiomes^20, 21^, aluminum tolerance^22^, defense responses^23^, and even inspiration for engineering applications^24^ (Figure S1).

**Fig. 1.**
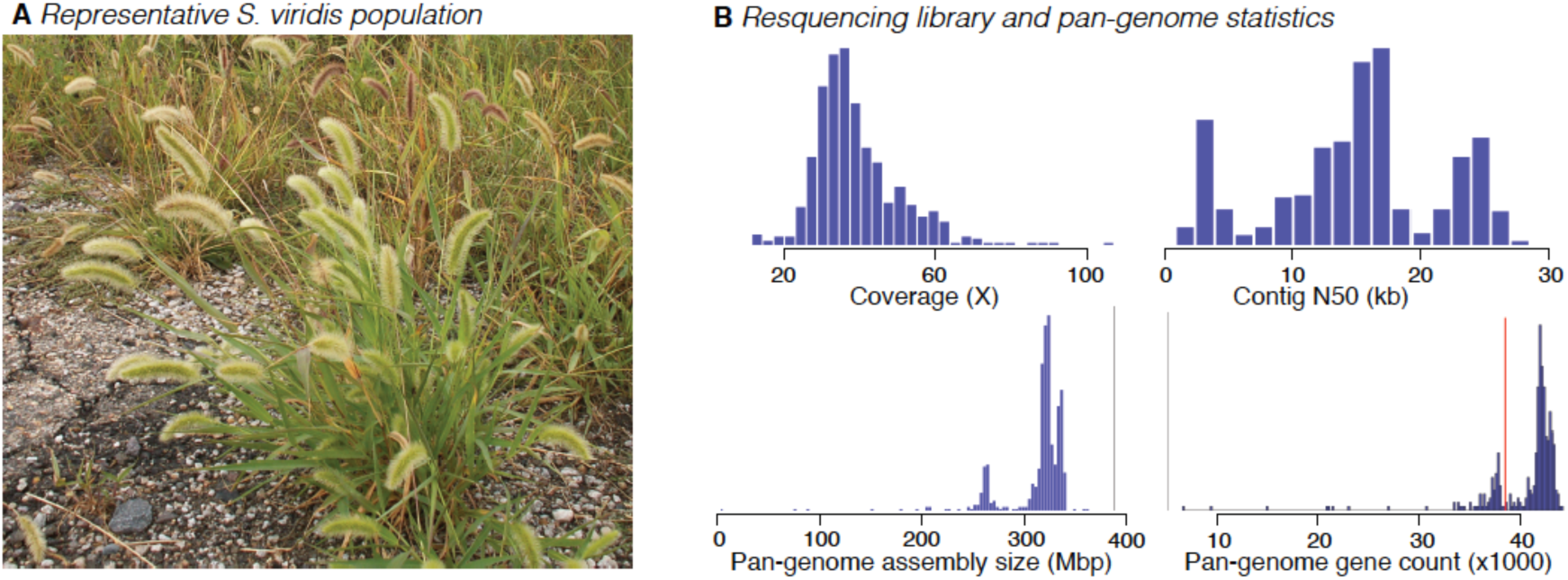
**a** *Setaria viridis*, in its common highly disturbed habitat next to a road. **b** Average library coverage, contig N50 (Kb), pan-genome assembly size, and number of genes per library; red vertical line in lower right panel represents the number of genes necessary for a library to be included for PAV analysis (n=39,000).

As in most wild species, seeds of *S. viridis* fall off the plant at maturity, a process known as shattering^25^. While essential for dispersal in natural ecosystems, shattering is undesirable in cultivation and humans have selected for non-shattering mutants since the dawn of agriculture^26–28^. Such domesticates include *Setaria italica* (foxtail millet), the domesticated form of *S. viridis*, which is grown as a crop in Asia^29–31^. Improvement of some current crops (e.g. African fonio and North American wildrice) is hindered because shattering causes high losses at harvest^32–34^. Thus identification of genes critical for shattering has direct agronomic benefit.

Human selection on wild grasses could only have been effective if there is standing variation in genetic loci that affect shattering. However, to our knowledge no previous study has succeeded in cloning such loci via association studies of natural diversity. Most previous efforts have instead relied on crosses between domesticated plants and their wild progenitors, and genes and QTL identified in such studies are, for the most part, not conserved among species^25, 35, 36^. We infer that many shattering-related loci remain to be identified.

In the work reported here, we surveyed natural populations of *S. viridis* and found allelic variation at a novel locus associated with seed shattering. We deployed a large sequence-based resource for *S. viridis*, analyzed its population structure using single-nucleotide polymorphisms (SNPs) and presence-absence variation (PAV) of individual genes, and tested for signatures of selection. Genome-wide association studies (GWAS) identified QTL for response to the abiotic environment and also a shattering-related locus, *Less shattering1* (*Les1*). Function of *Les1* was validated with genome editing. The orthologous gene in *S. italica* is disrupted by a transposon, indicating that the locus also contributed to domestication. In a parallel study, we identified a locus controlling leaf angle, a trait with clear agronomic implications. Data presented here show that genomics and biotechnological resources in *S. viridis* can be used to accelerate our mechanistic understanding of genetic processes and thereby contribute to enhanced and stabilized yield.

### A complete genome and genomic model for the Panicoideae

We have developed significant new resources for the *Setaria* community. A new assembly was generated for the *S. viridis* reference line ‘A10.1’, using a combination of long-read Pacbio and Illumina sequencing technologies. The final version 2.0 release is a complete telomere-to-telomere chromosomal assembly, containing 395.1 Mb of sequence in 75 contigs with a contig N50 of 11.2 Mb and a total of 99.95% of assembled bases in nine chromosomes. This is a major improvement over previous *S*. *viridis* genome releases, which had a contig N50 of 1.6 MB, and ca. 94% of reads in contigs. The associated gene annotation is equally complete (BUSCO score ^37^ on Embryophyta for the gene set v2.1 is 99%) describing 38,334 gene models and 14,125 alternative transcripts.

To complement the genome assembly and probe the genetic architecture of complex traits we conducted deep resequencing (mean of 56M high quality paired end reads, 42.6X coverage) of 598 *S. viridis* diversity samples using Illumina 2×150 paired end (PE) libraries (Figure 1b; metadata and sequence accession numbers in Table S1). Each library was subsequently assembled into a pan-genome database. While less contiguous than the long-read based A10.1 genome (mean n. contigs: 75,001; Contig N50: 16.2 Kb), total assembled bases were very similar (mean assembled bases = 322.5 M) (Figure 1b).

### Multiple introductions from Eurasia underlie distinct North American gene pools

The history of *S. viridis* in North America is unknown, although previous phylogenetic studies place it within a clade of Asian genotypes ^38, 39^; indicating that Eurasia is the native range while North America represents a recent and likely human-associated range expansion. To understand the relationship of the North American samples to samples from the Old World, we called 8.58M single nucleotide polymorphisms (SNPs) among our 598 Illumina libraries (Table S1). We then extracted polymorphisms comparable to the genotype-by-sequencing (GBS) data from previously-published Illumina sequence data for 89 non-US samples collected in China, Canada, Europe and Middle East ^40^. We then tested for population structure using fastStructure ^41^ and Admixture ^42^. Overall, we find four distinct subpopulations, all of which are found both in North American and Eurasia (Figure 2a, b). This pattern is expected if *S. viridis* diversified throughout Eurasia, was introduced to North America from several different sources, and then dispersed widely.

**Fig. 2.**
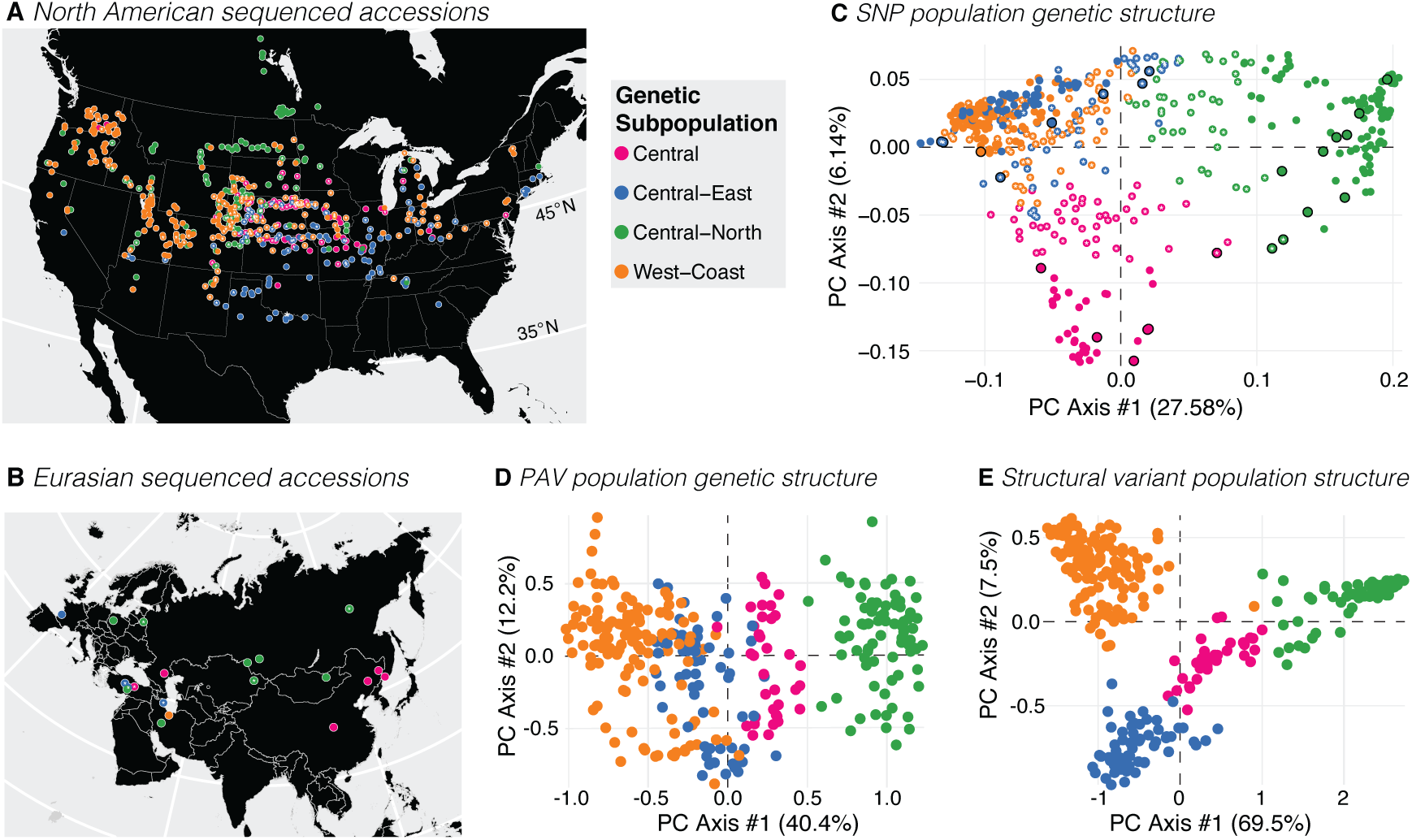
PAV and SNP diversity of subpopulations. **a** Geographic distribution and population assignment of North American accessions based on SNP data. **b** Geographic distribution and population assignment of Eurasian accessions based on SNP data. **c** PCA of SNP data, showing placement of North American samples among Eurasian ones. **d** PCA of PAV variants, excluding admixed individuals. **e** PCA of structural variants, excluding admixed individuals. Both SNP and PAV data clearly separate the Central-North population (green) from the others, and the structural variants distinguish all populations. *, accession is admixed; black rim on circle, accession is in its native range.

To define the evolutionary and biogeographical history of North American *S. viridis*, we conducted an identical population structure analysis with all SNPs in our resequencing panel. Of the 8.58 million SNPs (one SNP for every 21.6 bp), 430K mapped to exons (primary transcripts), 182K SNP were missense, 5K were nonsense, and 243K were silent. As with the GBS data, this analysis identified four main sub-populations (Central; Central-East; Central-North; West-coast; Figure S2). Not all individuals clustered uniquely into a single subpopulation; 382 genotypes were ‘pure’ and assigned to a single subpopulation while 216 individuals were ‘admixed’ (Q < 0.7). These subpopulations trace their origins to distinct Eurasian gene pools: the ‘Westcoast’ subpopulation was closely related to samples from a genetic subpopulation that spanned the northern Middle East region; the closest relatives of the USA ‘Central-North’ subpopulation were found from northern Europe through Siberia; the Eurasian members ‘Central-East’ subpopulation were restricted to northern Iran and Afghanistan; and samples from China exclusively group with the ‘Central’ USA subpopulation (Figure 2b, c). The Central USA population, whose Eurasian relatives inhabit a range from China to Turkey, had the highest SNP diversity, highest number of private alleles, and lowest mean linkage disequilibrium (LD) of all the populations (Table S2). Such elevated diversity could be driven by larger or multiple founding populations.

### Pan-Genome gene presence-absence variation mirrors SNP diversity

While SNPs serve as excellent proxies of total genetic diversity, previous work has demonstrated that larger-scale variation such as gene presence-absence and structural variants may underlie adaptation in nature and crop improvement^43–46^. To capture the set of all genes in all accessions of *S. viridis* (i.e., the pan-genome), we needed to identify genes that were missing or not annotated in the v2.1 gene set reference. Accordingly, we identified proteins in *Setaria italica* (v2.2), *Zea mays* (v.PH207) and *Sorghum bicolor* (v3.1) that were not shared with (i.e., not orthologous to) those in *S. viridis* (v2.1) and determined which were present in at least one member of the non-admixed diversity panel (382 accessions). This set of proteins was then added to the set from *S. viridis* (v2.1) to identify a pan-genome of 51,323 genes. Within this pan-genome, a core set of 39,950 genes occurred in 98% of all individuals. Of these core genes 32,732 were annotated in *S. viridis* (v2.1), with 3224, 3412, and 582 more identified based on similarity to *S. italica*, *Z. mays* and *S. bicolor* gene models, respectively. Discriminant function analysis of principal components (DAPC)^47, 48^, a multivariate method for identifying clusters of genes, identified an additional 5385 genes in 56% of all individuals (the “shell” set), and found the remaining 5987 in 12% of all individuals (the “cloud”).

Consistent with other studies of pan-genome population genetics^49^, the SNP- and PAV-based estimates of subpopulation structure are similar. Analysis of the “shell” gene PAV data revealed four subpopulations (n = 130, 78, 59, 35, respectively; Fig. 2d). Of the 203 non-admixed genotypes (that also had sufficient PAV data), 190 (94%) assigned to the same genetic subpopulation as in the SNP analysis. The Central-North population is clearly distinct in both PAV and SNP data (Fig. 2c), and when all structural variants are considered, the four populations are distinct (Fig. 2e). 4,062 genes are significantly over- or under-represented among subpopulations (*P* < 0.05; X^2^-test, with Benjamini-Hochberg correction), with 45 private alleles (MAF > 0.1) specific to particular subpopulations (Table S3). KEGG and GO enrichment analyses for over-represented genes in each population found pathways relating to biosynthesis of secondary compounds, and genes involved in defense response to pathogens or herbivores Figure S3.

Our PAV data connect published QTL studies with population-level processes. For example, a study of drought response in *S. viridis* found a strong-effect QTL significant for plant size, water loss, and WUE across all tested environments ^50^. Investigation within the pan-genome found an *S-phase kinase associated protein1* (*Skp1)* homolog (Sevir.2G407800) within 50Kb of the associated SNP that was common in the Central-North population (present in 65% of individuals), while significantly under-represented in the Central-East (present in 5% of individuals) (p <4.3 × 10^−15^). SKP1 proteins are part of the E3 ubiquitin ligase complex that leads to protein degradation^51^; while the physiological role of this particular SKP1 homolog is unknown, the fact that it falls within a known QTL interval and is differentially represented in two populations suggests it may be a candidate for future investigation.

### Selection and correlation with climate in the new range of *S. viridis*

Combined, the PAV and SNP data clearly demonstrate massive genetic diversity and distinct gene pools to target molecular dissection of agriculturally important traits, including response to the environment. While we have shown that genetic variation in North American *S. viridis* reflects multiple introductions, we also observe pervasive genetic and biogeographic admixture. These factors, combined with its rapid cycling and weedy life history, indicate that adaptation to climate may underscore a biogeographic range expansion across much of the continent. To test this hypothesis, we characterized the environment for each accession using principal components analysis (PCA) of the Worldclim^52^ environmental variables. To overcome collinearity among the climate variables (Figure S4), we transformed the 19 variables into the first three principal components (36.13%, 30.07% and 16.36% variance explained; factor loadings in Table S4), which served as response variables for three GWAS analyses. To control for population structure, we also supplied a set of three SNP-derived kinship PC axes which explain 15.34% of the genetic relatedness in our panel.

While the first bioclimatic PC axis was not significantly associated with any SNP markers, PC2 had one association (Chr01: 37,770,697) (Figure S5) and PC3 identified several peaks with a total of 140 significant markers (Bonferroni corrected p-value, 5.82e-09). PC3 is loaded by climatic variables relating to extremes of precipitation and temperature. Of the 140 PC3 hits, 66 fall within 16 genes (Table S5). Genes within 100 Kbp of the significant markers were associated with organellar genome maintenance (GO:0033259, GO:0000002, GO:0006850), protein breakdown (GO:0045732), gibberellic acid homeostasis (GO:0010336), and fucose metabolism (GO:0006004, GO:0042353) (Table S6).

Despite the relatively small number of clear associations with parameters of the abiotic environment, both Tajima’s D and iHS found multiple genes under selection in the four populations (Figure S6; Table S7). The two tests have different underlying assumptions and methods; genes that appear as outliers in both tests and annotations that arise repeatedly are excellent candidates for further investigation. For example, in most subpopulations we find selected genes consistently enriched in flavonoid (and other derivatives of phenylalanine) metabolism GO terms and KEGG pathways, processes that are often underlie responses to herbivores and pathogens and could be involved in local adaptation. However, selection on other processes and pathways appears to be specific to only one subpopulation. For example, genes involved in pH reduction in the Central-North population are identified by both tests, although interpretation of this result would require investigation of individual sets of genes and their tissue localization. Together, the bioclim GWAS and tests for selection show that *S. viridis* can provide testable hypotheses of gene function and phenotypic output.

### A novel gene, *SvLes1*, controls seed shattering in *S. viridis*

We deployed the new high quality genome, pan-genome and population genetic analyses to link genotypic diversity to the agronomically important phenotype of reduced shattering. We tested a subset of lines for seed shattering using a simple shattering index, in which mature panicles were scored for shattering on a scale of 1 (low) to 7 (high; Table S8). GWAS identified a single strong QTL (peak -log_10_ P > 30) for seed shattering on chromosome 5 (Figure 3a) a region of approximately 2Mb above the experiment-wise p=0.01 Bonferroni correction threshold. In this region, 119 mutations, primarily missense, alter protein sequences relative to the reference; we used PROVEAN^53^ to predict deleterious mutations that are more likely to alter the biological function of protein products. Combining this prediction and the association score of SNPs (Figure 3b), we prioritized a single G-T polymorphism (Chr_05:6849363) in a gene encoding a MYB transcription factor, *SvLes1* (Sevir.5G085400, similar to Sobic.003B087600 from *Sorghum bicolor* and ZM00001d040019_T001 from *Zea mays*), with two MYB DNA-binding domains. The mutation leads to a R84S substitution in the second MYB domain of *SvLes1* (Figure 3c). We name these two alleles *SvLes1-1* and *SvLes1-2*, associated with high seed shattering and reduced shattering, respectively. The *SvLes1-2* allele appears in 24% of the 215 accessions of the GWAS panel, and clearly associates with reduced shattering scores (p=6.53E1-33, 2-tailed test; Table S8). The reference line A10.1 is among these reduced-shattering lines.

**Fig. 3.**
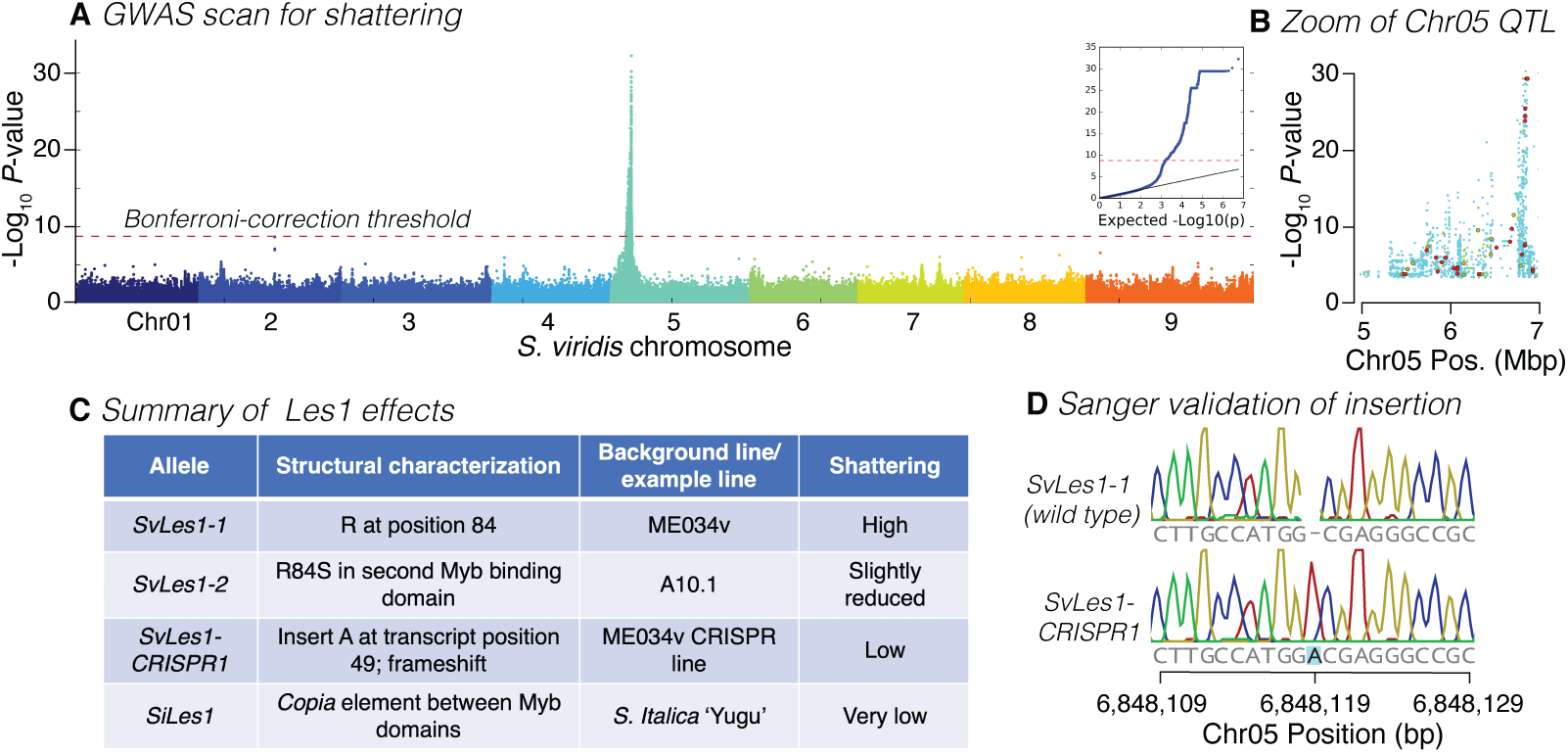
GWAS and cloning of *Les1*. **a** Manhattan plot of GWAS result, red line showing p=0.01 after Bonferroni correction. **b** Zoom in to peak on Chromosome 5; larger dots represent missense SNPs identified by snpEff. Different colors of missense SNPs indicate Provean score range (blue for >−2.5, green for <−2.5 and >−4.1, red for <−4.1; −2.5 and −4.1 represent 80% and 90% specificity). Lower scores indicate higher likelihood of deleterious effects of the mutation. **c** Table of *Les1* alleles in *S. viridis* (*Sv*) and *S. italica* (*Si*), with structural characteristics, background line, and shattering phenotype. **d** Sanger sequence validating the position of the adenine insertion (frameshift) in *SvLes1-CRISPR-1*.

To validate *SvLes1* as the causal gene, we used CRISPR-Cas9 to create additional alleles. We disrupted the wild type, high-shattering allele *SvLes1-1* in the accession ME034v (corresponding to accession TB0147) to create several novel, non-functional alleles. Sequence analysis of *SvLes1-CRISPR1* revealed an adenine insertion at position 149 of the transcript (Figure 3d), leading to a frameshift mutation predicted to completely abolish gene function thereby creating non-shattering plants. After segregating out the transgenes encoding Cas9 and guide RNAs, homozygotes of *SvLes1-CRISPR1* were phenotypically examined in the T3 generation.

To quantify seed shattering, we measured tensile strength of the abscission zone (AZ)^54, 55^. We compared *SvLes1-CRISPR1* and *SvLes1-1*(both in an ME034v background) with *SvLes1-2* (in the A10.1, reduced shattering, background). *SvLes1-1* had the lowest tensile strength (high shattering), *SvLes1-2* tensile strength was slightly higher (less shattering), and *SvLes1-CRISPR1* had high tensile strength (reduced seed shattering) (Figure 4a,b). These results were confirmed with wind tunnel experiments measuring the number of seeds released from the inflorescence and the distance they traveled (Figure 4c). Few *SvLes1-CRISPR1* seeds were released from the plant, whereas dozens of seeds were released from the *SvLes1-1* plants at six weeks after heading. Seeds of the *SvLes1-CRISPR1* allele weighed significantly less than in *SvLes1-1*, but germination percentage was unaffected (Figure S7). T3 offspring of T2 heterozygous plants segregated 3:1 shattering to non-shattering as expected for an induced mutation in a single gene, implying that the lines are isogenic except for the mutant allele.

**Fig. 4.**
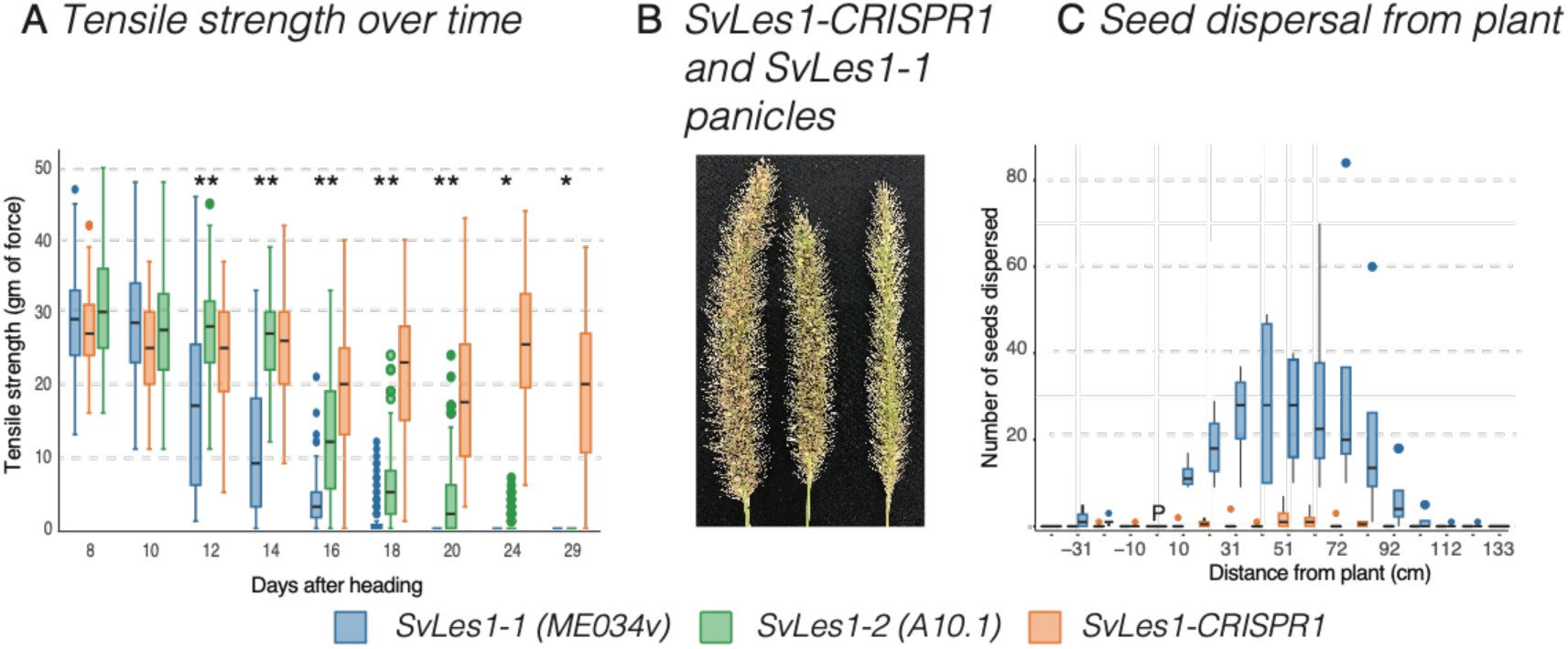
Phenotypic characterization of *SvLes1-CRISPR1* mutants and naturally occurring alleles. **a** Force required to break the AZ (tensile strength in gm of force) in *SvLes1-1*, *SvLes1-2*, and *SvLes1-CRISPR1* lines, measured every two days starting at 8 days after heading. **b** Image of *S. viridis* high shattering (*SvLes1-1* in ME034), left two panicles) and *SvLes1-CRISPR1* mutant panicles seven weeks after heading. Both mutants have retained their seed, unlike ME034 which has shed most of its seed at this time point. **c** Seed dispersal distances in a wind tunnel, measured from four independent high shattering plants (*SvLes1-1* in ME034) and eight *SvLes1-CRISPR1* plants at week 6. Wind speed was 5 mph coming from the left of the diagram. The plant, indicated by P, was placed in middle of the tunnel; some seed fell on the windward side (negative distance values) due to oscillation of the plant. Although mean dispersal distance was approximately the same for both alleles, the *SvLes1-CRISPR1* plants released significantly fewer seeds at each distance that the *SvLes1-1* plants.

*SvLes1* is a transcription factor that has not been implicated in shattering in any other species. Despite studies identifying shattering related genes in rice (e.g. *Sh4*^55^, *qSh1*^56^, *Shat1*^57^) and sorghum (e.g. *Sh1*^58^), the cellular mechanism of shattering is not known in any cereal, and recent data suggest that each species may be unique^36^. Unlike other grasses, the AZ in *Setaria viridis* is not histologically distinct and is only subtly different from that in *S. italica*^25, 54^. Cells and cell walls in the AZ are not clearly different from their neighboring cells (demonstrated by multiple different cell wall stains plus TEM^36^). Given this histology, we expected that the AZ of non-shattering mutants would look like that of wild type plants, and indeed the anatomy and histology of *SvLes1-CRISPR1* and *SvLes1-1*spikelets are indistinguishable (Figure S8).

### Recent transposable element insertion in *S. italica Les1* contributed to domestication of foxtail millet

Because *Setaria italica* (foxtail millet) is a domesticated (non-shattering) derivative of *S. viridis*^29–31^, we hypothesized that selection of non-shattering lines could have identified rare alleles of *S. viridis* or *S. italica* early in the domestication process. We discovered a ∼6.5kb *copia* transposable element (*copia38*) inserted between the two Myb domains of *SiLes1* in the *S. italica* line Yugu1. We call this the *SiLes1-TE* (transposable element) allele. The disruption of the Myb domain strongly suggests *SiLes1-TE* is a loss of function allele similar to *SvLes1-CRISPR1* and should also produce a low shattering phenotype, thus potentially contributing to the domestication of foxtail millet. The *copia38* transposable element was aligned to each of the 598 *S. viridis* re-sequenced lines to investigate whether any had the TE insertion in *SvLes1*. Only two samples aligned to the *copia38* TE within the CDS sequence of *SvLes1*, but the nucleotide identity and coverage of the alignments were poor (32% and 6% respectively). In contrast, *Copia38* is nearly fixed among the foxtail millet lines examined (78 out of 79).

Genome-wide, *S. italica* has about 22% of the SNP variation of *S. viridis* (based on sequences generated in ^59^), as expected given its domestication history. In the *SiLes1* region, however, this number is significantly reduced to 4.1-8.2% of the diversity in *S. viridis* depending on the size of the region compared (10-100 Kbp, Table S9) (p=0.0066 based on 100,000 coalescent simulations of the ratio of π *italica*/π *viridis*). These data hint that the low-shattering QTL might co-localize with a selective sweep. Diversity is relatively high within the *S. italica* gene itself, which may mean that selection is on a regulatory region or additional locus under the QTL or that the transposon insertion has rendered *SiLes1* a pseudogene. Estimates of linkage disequilibrium (LD) support this interpretation (Table S10). In *S. viridis* estimates of LD do not vary much across intervals from 10 to 100 Kbp surrounding *SvLes1*. In *S. italica* on the other hand, LD is nearly complete for the 20 kb region surrounding *SiLes1*. When that region is expanded to 40 Kbp LD drops to levels approximating that in *S*. *viridis*. Taken together, these data indicate either a selective sweep or purifying selection on a genomic region that includes *SiLes1* and *copia38* in LD.

*SiLes1* has not been identified in other studies of *S. italica* and thus its potential role in domestication is newly described here. Two strong QTL for shattering were identified in recombinant inbred lines derived from an *S. italica* x *S. viridis* cross (accessions B100 and A10.1, respectively)^60^, but neither QTL encompasses *SiLes1*. Analysis of 916 diverse accessions of *S. italica* identified 36 selective sweeps, but the *SiLes1* region does not colocalize with any of them^59^. In addition, a previous study^54^ found that tensile strength in two elite *S. italica* lines, Yugu1 and B100, is higher than that of the *SvLes1-CRISPR1* homozygotes reported here. Thus all previous data indicate that *SiLes1-TE* is but one of several loci contributing to the lack of shattering in foxtail millet.

*Copia38* is a long terminal repeat (LTR) retroelement with two 451 bp LTR sequences. The LTRs for *Copia38* are identical across the entire pan-genome, suggesting that the insertion is recent, because mutations start to accumulate in the once-identical LTRs immediately after insertion of a TE. Phylogenetic analysis showed *Copia38* tightly clusters with a few homologous copies on a long branch, indicating a shared burst event for copies in this cluster (Figure S9). Pairwise distance among the copies suggests this burst was recent, on average about 45k (range 23k – 81k) years ago, assuming a neutral mutation rate of 6.5×10^−9^/bp/year^61^. The copies from this burst only occur in Yugu1 but not in A10.1, suggesting a recent expansion just prior to domestication of *S. italica* that began approximately 8000 years ago^29–31^.

Combining the results and evidence above, a plausible scenario is that the *SiLes1-TE* allele was a low frequency allele that was selected during domestication due to its favored low shattering phenotype, and spread quickly through the early foxtail millet land races. Later, the low shattering phenotype was further strengthened by additional loci with stronger effects (i.e. *SvSh1*^59, 60^) during recent crop improvement. Genome editing technology today allows us to recreate a low-shattering phenotype from ancestral *S. viridis* alleles, mimicking the initial phase of foxtail millet domestication through a different novel allele *SvLes1-CRISPR1*.

### Parallel genetic control of leaf angle in *S. viridis* and maize by *liguleless2* orthologs

The diversity panel is also a powerful tool for uncovering the basis of rare traits, including those of agronomic value. We discovered a single accession in the panel (TB0159) with reduced auricle development and markedly upright leaves (small leaf angle) (Figure 5a,b), a trait of considerable interest in the major crops maize and rice, where it has led to increased planting densities and higher yields^62–64^. As GWAS is not suitable for mapping traits with low frequency and strong effects, bulked segregant analysis (BSA)^17, 40^ was used to identify the associated genomic region. BSA is a process in which plants with and without a particular phenotype are pooled (bulked) and then each pool is sequenced. Regions or loci that differ between the two pools are then inferred to contain the mutations underlying the phenotype. TB0159 was crossed to A10.1, and the F1 plants showed wild type leaf angle, showing that small leaf angle is recessive. The wild type and small leaf angle trait in the F_2_ population segregated at 264:153 which differs significantly from a 3:1 ratio (p=0.000238). This could be explained by partial dominance at a single locus, or by several loci controlling the phenotype. We proceeded assuming a single partially dominant causal gene for small leaf angle.

**Fig. 5.**
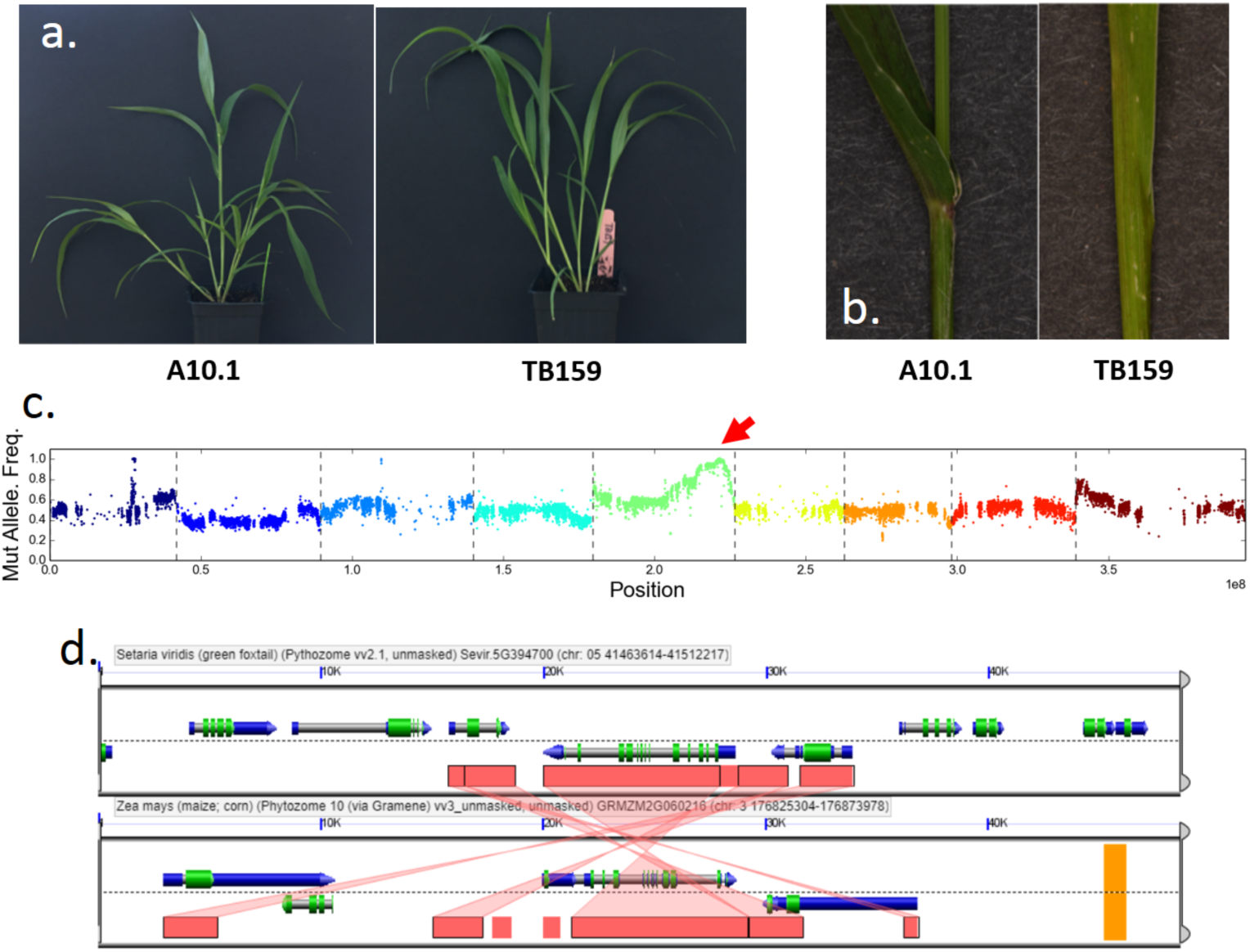
Phenotype and mapping of small leaf angle. **a, b** Small leaf angle phenotype in TB159. **c** BSA mapping result, red arrow indicating QTL. **d** Synteny analysis around *SvLg2* and maize *lg2* locus.

With BSA, we coarsely mapped the reduced leaf angle phenotype to a homozygous region of ∼800 kb on chromosome 5 (Figure 5c) that contained 104 disruptive SNPs and 687 indels (393 single bp). This region includes *SvLiguless2* (*SvLg2*) (Sevir.5G394700), the syntenic ortholog of *liguless2* in maize (Figure 5d), which is a transcriptional regulator that controls auricle development and leaf angle^65^. Because the small leaf angle phenotype is partially dominant and unique to TB0159 in the panel, the causal allele should be homozygous and occur only in that accession. We identified a homozygous G insertion (Chr_5:41489494) in the coding region of *SvLg2* of that is predicted to cause a frameshift. Accordingly, we hypothesize that this indel mutation might be the causal mutation for the small leaf angle in TB0159. As the ligule is a trait unique to grasses and has been a target for breeding programs, *S. viridis* can now be used as a model system to rapidly dissect the molecular genetic network of Lg2, which remains poorly resolved in maize and rice, to identify additional candidates for breeding or engineering of this important architectural trait.

In summary, data presented here demonstrate the power of the *S. viridis* genome and associated diversity lines for gene discovery. Together these tools permit discovery of novel environmental associations and genes such as *SvLes1*, and additional insight into genes such as *SvLg2* that have been identified from related systems. We show that *S. viridis* resources can uncover the genetic basis of a trait when the genes are unknown and likely differ from those in other related species. We also show that *S. viridis* can be a model for investigating genes in which an orthologue is known in a crop. Specifically, we have used the *S. viridis* genome and diversity panel to study the genetic basis of seed shattering and leaf angle (*SvLes1* and *SvLg2*, respectively), traits that influence crop yield by affecting harvestability, productivity and density of planting.

## Methods

### Plant materials

The reference line A10.1 is a descendant of the line used by Wang et al.^66^ in early RFLP maps. The original line was found to be heterozygous and thus A10.1 was propagated via single-seed descent by Andrew Doust (Oklahoma State University, Stillwater, OK, pers. comm.). It is thought to have originated in Canada. The other reference, ME034 (also known as ME034v), was collected by Matt Estep (Appalachian State University, Boone, NC) in southern Canada as part of a diversity panel^39^ included among the diversity lines sequenced here. Transformation is more efficient for ME034 than for A10.1 (Joyce van Eck, Boyce Thompson Institute, Ithaca, NY, pers. comm.) and thus the former is being used widely for functional genetic studies.

The 598 individuals of the diversity panel were collected over a period of several years. About 200 lines have been described in previous studies^39, 67^ whereas others were new collections added for this project. Individuals were propagated by single seed descent, although the number of generations varies by accession. *Setaria viridis* is inbreeding by nature (ca. 1%^67^), so we assume that initial heterozygosity was generally low and then further reduced in propagation.

### Library creation and sequencing

To prepare DNA for sequencing of the reference line, 100 ng of DNA was sheared to 500 bp using the Covaris LE220 (Covaris) and size selected using SPRI beads (Beckman Coulter). The fragments were treated with end-repair, A-tailing, and ligation of Illumina compatible adapters (IDT, Inc) using the KAPA-Illumina library creation kit (KAPA biosystems). The prepared library was then quantified using KAPA Biosystem’s next-generation sequencing library qPCR kit and run on a Roche LightCycler 480 real-time PCR instrument. The quantified library was then multiplexed with other libraries, and the pool of libraries was prepared for sequencing on the Illumina HiSeq sequencing platform using a TruSeq paired-end cluster kit, v3, and Illumina’s cBot instrument to generate a clustered flowcell for sequencing. Sequencing of the flowcell was performed on the Illumina HiSeq2000 sequencer using a TruSeq SBS sequencing kit, v3, following a 2×150 indexed run recipe.

Plate-based DNA library preparation for Illumina sequencing was performed on the PerkinElmer Sciclone NGS robotic liquid handling system using Kapa Biosystems library preparation kit. 200 ng of sample DNA was sheared to 600 bp using a Covaris LE220 focused-ultrasonicator. Sheared DNA fragments were size selected by double-SPRI and then the selected fragments were end-repaired, A-tailed, and ligated with Illumina compatible sequencing adaptors from IDT containing a unique molecular index barcode for each sample library.

The prepared library was quantified using Kapa Biosystem’s next-generation sequencing library qPCR kit and run on a Roche LightCycler 480 real-time PCR instrument. The quantified library was then multiplexed with other libraries, and the pool of libraries was then prepared for sequencing on the Illumina HiSeq sequencing platform using a TruSeq paired-end cluster kit, v3 or v4, and Illumina’s cBot instrument to generate a clustered flowcell for sequencing. Sequencing of the flowcell was performed on the Illumina HiSeq2000 or HiSeq2500 sequencer using HiSeq TruSeq SBS sequencing kits, v3 or v4, following a 2×150 indexed run recipe.

### Sequencing of the reference genome

We sequenced *S. viridis* A10.1 using a whole genome shotgun sequencing strategy and standard sequencing protocols. Sequencing reads were collected using Illumina HISeq and PACBIO SEQUEL platforms at the Department of Energy (DOE) Joint Genome Institute (JGI) in Walnut Creek, California and the HudsonAlpha Institute in Huntsville, Alabama. One 800 bp insert 2×250 Illumina fragment library (240x) was sequenced, giving 425,635,116 reads (Table S11). Illumina reads were screened for mitochondria, chloroplast, and PhiX contamination. Reads composed of >95% simple sequence were removed. Illumina reads <75bp after trimming for adapter and quality (q<20) were removed. For the PACBIO sequencing, a total of 36 P5C2 chips (4 hour movie time) and 41 P6C4 chips (10 hour movie time) were sequenced with a p-read yield of 59.09 Gb, with a total coverage of 118.18x (Tables S11, S12).

### Genome assembly and construction of pseudomolecule chromosomes

An improved version 2.0 assembly was generated by assembling 4,768,857 PACBIO reads (118.18x sequence coverage) with the MECAT assembler^68^ and subsequently polished using QUIVER^69^. The 425,635,116 Illumina sequence reads (240x sequence coverage) were used for correcting homozygous snp/indel errors in the consensus. This produced 110 scaffolds (110 contigs), with a contig N50 of 16.8 Mb, and a total genome size of 397.9 Mb (Table S13). A set of 36,061 syntenic markers derived from the version 2.2 *Setaria italica* release was aligned to the MECAT assembly. Misjoins were characterized as a discontinuity in the *italica* linkage group. A total of 15 breaks were identified and made. The *viridis* scaffolds were then oriented, ordered, and joined together into 9 chromosomes using syntenic markers. A total of 61 joins were made during this process. Each chromosome join is padded with 10,000 Ns. Significant telomeric sequence was identified using the TTTAGGG repeat, and care was taken to make sure that it was properly oriented in the production assembly.

Scaffolds that were not anchored in a chromosome were classified into bins depending on sequence content. Contamination was identified using blastn against the NCBI nucleotide collection (NR/NT) and blastx using a set of known microbial proteins. Additional scaffolds were classified as repetitive (>95% masked with 24mers that occur more than 4 times in the genome) (26 scaffolds, 1.2 Mb), alternative haplotypes (unanchored sequence with >95% identity and >95% coverage within a chromosome) (15 scaffolds, 1.0 Mb), chloroplast (3 scaffolds, 164.5 Kb), mitochondria (5 scaffolds, 344.9 Kb), and low quality (>50% unpolished bases post polishing, 1 scaffold, 19.3 Kb). Resulting final statistics are shown in Table S14.

Finally, homozygous SNPs and INDELs were corrected in the release consensus sequence using ∼60x of Illumina reads (2×250, 800 bp insert) by aligning the reads using bwa mem ^70^ and identifying homozygous SNPs and INDELs with the GATK’s UnifiedGenotyper tool ^71^. A total of 96 homozygous SNPs and 4,606 homozygous INDELs were corrected in the release. The final version 2.0 release contains 395.1 Mb of sequence, consisting of 75 contigs with a contig N50 of 11.2 Mb and a total of 99.95% of assembled bases in chromosomes (Table S14).

Completeness of the v2.0 assembly is very high. To further assess completeness of the euchromatic portion of the version 2.0 assembly, a set of 40,603 annotated genes from the *S. italica* release was used for comparison. The aim of this analysis is to obtain a measure of completeness of the assembly, rather than a comprehensive examination of gene space. The transcripts were aligned to the assembly using BLAT ^72^ and alignments ≥90% base pair identity and ≥85% coverage were retained. The screened alignments indicate that 39,441 (97.14%) of the *italica* genes aligned to the version 2.1 release. Of the unaligned 1,162 transcripts, 928 (2.28%) indicated a partial alignment, and 234 (0.58%) were not found in the version 2.1 release.

To assess the accuracy of the assembly, a set of 335 contiguous Illumina clones >20 Kb was selected. A range of variants was detected in the comparison of the clones and the assembly. In 239 of the clones, the alignments were of high quality (< 0.01% bp error) with an example being given in Figure S10a (all dot plots were generated using Gepard ^73^). The remaining 96 clones indicate a higher error rate due mainly to their placement in more repetitive regions (Figure S10b). The major component of the error in the 96 repetitive clones was copy number variation, which affected 50 of the clones. These 50 clones accounted for >97% of all of the errors in the 335-clone set. Excluding the clones with copy number variation, the overall bp error rate in the 285 clone set is 0.0098% (1,043 discrepant bp out of 10,601,785).

### Annotation

Various Illumina RNA-seq reads were used to construct transcript assemblies using a genome-guided assembler, PERTRAN^74^: 1B pairs of geneAtlas, 0.9B pairs of LDHH, 0.9B pairs of LLHC, and 176M other pairs. 109,119 transcript assemblies were constructed using PASA^75^ from RNA-seq transcript assemblies above.

Gene models of v1.1 on assembly v1.0 were lifted over to assembly v2.0 and improved. The in-house gene model improvement (GMI) pipeline is as follows:

The genomic sequence of a locus is obtained, including introns if any and up to 1Kbp extensions on both ends unless they run into another gene. For intergenic space less than 2Kbp, half of the intergenic distance is the extension for 2 adjacent loci. These locus sequences are mapped to a new genome using BLAT. Duplicate mappings are resolved using the gene model’s neighbors in original genome space. When a locus genomic sequence is mapped to the new genome uniquely and 100%, the gene model is perfectly transferred to the new genome. For the remaining gene models, both their transcript and CDS sequences are mapped with BLAT to the region in the new genome located by locus genomic sequence mapping above. Gene models are made from CDS alignments with quality of 95% identity, 90% coverage and valid splice sites if any, and are transferred if the resulting peptide is 70% or more similar to the original gene model peptide. UTRs are added if any using transcript alignments. Remaining gene models are mapped to the new genome using GMAP. Gene models based on GMAP alignments with quality of 95% identity, 70% coverage and valid splice sites if any are transferred if and only if the resulting gene model peptide is 70% or more similar to the original gene model peptides and in the new genome location not occupied by transferred gene models in earlier steps.

Non-overlapping complete ORFs from each PASA transcript assembly (TA) were predicted if the ORF had good homology support or was long enough (300bp if multi exons or 500bp if single exon). Proteins from *Arabidopsis thaliana*, rice, sorghum, *Brachypodium distachyon*, *Setaria italica*, grape, soybean and Swiss-Prot eukaryote were used to score TA ORFs using BLASTP. The TA ORFs were then fed into the PASA pipeline where EST assemblies were obtained for gene model improvement including adding UTRs. PASA improved gene model transcripts were compared to v1.1 lifted over models on how well the transcript CDS was supported by ESTs and/or homologous protein, and not overlapped with repeats generated with RepeatMasker^76^ for more than 20 percent. If PASA gene models of TA ORFs were better than lifted over ones, the PASA gene models took over the lifted over ones. Otherwise, the lifted over gene model stayed. The final gene model proteins were assigned PFAM, PANTHER and gene models were further filtered for those with 30% or more of proteins assigned to transposable element domains.

Locus model name was assigned by mapping forward v1.1 locus model if possible using our locus name map pipeline; otherwise, a new name was given using JGI locus naming convention that was used in v1.1 locus model naming. Our locus name map pipeline is as following: a locus is said to be mapped and name mapped forward if 1) the prior version and current version loci overlap uniquely and appear on the same strand, and 2) at least one pair of translated transcripts from the old and new loci are MBH’s (mutual best hits) with at least 70% normalized identity in a BLASTP alignment (normalized identity defined as the number of identical residues divided by the longer sequence). For a given pair of prior version and current version transcripts at mapped loci, transcript model names are mapped forward if either a) an MBH relationship exists between the two proteins with at least 90% normalized identity or, b) the proteins have at least 90% normalized identity and are not MBH, but the corresponding transcripts sequences are (also with 90% normalized identity). This latter rule is specifically to handle cases where the prior version and current version models differ mainly by the addition of, or extension, of the UTR to a prior version model. These rules allowed the model names of approximately 92% v1.1 gene models mapped forward to v2.1.

### Sequencing and assembly of the diversity panel

After excluding seven lines because of low sequence coverage, 598 diversity samples (metadata, including Sequence Read Archive (SRA) numbers in Table S1) were used for diversity analysis. The samples were sequenced using Illumina paired end sequencing (2*151bp) at the Department of Energy Joint Genome Institute (JGI, Walnut Creek, CA, USA) and the HudsonAlpha Institute for Biotechnology (Huntsville AL, USA) using Hiseq 2500 and NovoSeq6000. Individual de novo assemblies for each line were constructed using Meraculous (v2.2.5)^77^ with a kmer size of 51, selected to maximize the contig N50 in the resultant assemblies, and to ensure that alternative haplotypes would have the best chance of being split apart. To construct chromosomes for each library, exons from the *S. viridis* gene set reference (v2.1; number of genes = 38,334; number of exons = 289,357) were aligned to each Meraculous assembly (blastn, word_size = 32), and exon alignments with identity >=90% and coverage >=85% were retained. Scaffolds were joined into gene-based scaffolds based on exon alignments; synteny and exon alignments were then used to order and orient the sequences into chromosomes (Figure S11).

### SNP calling

Reads from the diversity samples were mapped to *S. viridis* v2.1 using bwa-mem^78^. The bam was filtered for duplicates using picard (http://broadinstitute.github.io/picard), and realigned around indels using GATK^71^. Multi sample SNP calling was done using SAMtools mpileup^79^ and Varscan V2.4.0^80^ with a minimum coverage of eight and a minimum alternate allele frequency of four. An allele is confirmed to be homozygous or heterozygous using a binomial test for significance at a p-value of 0.05. Repeat content of the genome was masked using 24bp kmers. Kmers that occur at a high frequency, up to 5%, were masked. SNP around 25bp of the mask were removed for further analysis. A SNP was included for further analysis if it had a coverage in 90% of the samples, and a MAF > 0.01. Imputation and phasing were done in Beagle V4.0^81^. SNP Annotation was performed using snpEff^82^.

### Pan-genome and PAV analysis

To assess presence/absence variation (PAV) of genes across the diversity panel, admixed individuals were removed from the analysis, leaving 382 individuals (Q>=0.7). We expected that some genes present in wild accessions of *S. viridis* (i.e. the pan-genome) would be either missing or not annotated in the v2.1 reference gene set. To capture these, we included not only proteins from *S. viridis* (v2.1 gene set) but also non-orthologous proteins from *Setaria italica* (v2.2), *Z. mays* (v.PH207) and *S.bicolor* (v3.1)(based on InParanoid comparisons^83^) and aligned these to chromosome integrated assemblies from each of the four subpopulations using blat (-noHead -extendThroughN -q=prot -t=dnax).

Genes from *S. viridis* and *S. italica* were considered present if they aligned with greater than 85% coverage and identity, or at least 90% coverage and identity if the exons were broken up and located on no more than three contigs. *Sorghum bicolor* and *Zea mays* genes were considered present if they aligned with greater than 70% identity and 75% coverage (to allow for greater divergence among sequences), or at least 80% identify and coverage if the exons were broken up and located on no more than three contigs. Libraries with fewer than 39,000 genes considered present were excluded as genes were likely lost due to low coverage/poor assembly, with 302 individuals remaining for analysis. 67,079 genes were aligned to each assembly. After removing genes that aligned poorly or not at all to any assembly, a total of 51,323 genes was retained. The resultant PAV matrix (Table S15) was used to determine and cluster genes into their pan-genome designation (core, shell, cloud)^74–76^, using discriminant function analysis of principal components (DAPC)^41^. Using successive Kmeans clustering, three distinct clusters (based on BIC criteria) were discovered based on their PAV observance across non-admixed individuals, designated core, shell and cloud. Genomic coordinates for non-orthologous proteins in the pan-genome were determined using the GENESPACE pipeline^74^.

To discover which genes were over or under represented within each subpopulation based on their expected observance, a X^2^-test (with Benjamini-Hochberg correction) was performed for each gene among each of the four subpopulations. Significantly over-represented genes (allowing overlap among subpopulations; p<0.05) within subpopulations were also tested.

To infer syntenic regions when placing non-orthologous genes of interest back on the *S. viridis* genome, we applied the GENESPACE pipeline^74^ to the *S. viridis* genome described herein and five other grasses: *S. italica* (v2.2), *S. bicolor* (v3.1), *O. sativa* (‘Kitaake’, v3.1), *Z. mays* (‘Ensemble-18’) and *B. distachyon* (v3.1). Genome annotations and assemblies were downloaded from phytozome (phytozome.jgi.doe.gov). In short, GENESPACE runs the following pipeline. First it parses blast results to the top n (default = 1 * (ploidy / 2)) hits for each gene. A separate orthofinder run is conducted for each pairwise combination of genomes; only blast hits within orthogroups are retained. Syntenic blocks are formed with MCScanX^84^ with the block size (-s) and gap (-m) parameters set to 25 and 50 respectively. Blocks are culled to those with high mapping density through 2-dimensional nearest neighbor clustering implemented in the R package dbscan^85^. The resulting pruned collinear blast hits are used as anchors for fixed radius nearest neighbor search based on gene rank. All blast hits within the radius (default r = 100 genes) are considered syntenic. Syntenic blast from all species comparisons are fed into a final orthofinder run and parsed into orthologs and paralogs. Plotting routines are implemented in base R.

### Structural Variants

To detect structural variants within the pan-genome, pseudo PacBio reads were generated from assemblies of non-admixed individuals. Pseudo-reads (length: 10kb; depth 5X) were generated from all contigs greater than 10Kb from each pan-genome assembly. The pseudo-reads were aligned to the *S. viridis* reference genome using nglmr (v0.2.7)^86^ with default settings for Pacbio reads. The resulting bam file was sorted using samtools (v1.10) and used for calling structural variants with sniffles (v1.0.11). The structural variant types considered across the pan-genome were: insertions, deletions, and inversions. The average number of SV’s detected per library was 15,593. A presence/absence matrix for each SV type and was clustered using iterative Kmeans and BIC to determine goodness of fit and the optimal number of clusters present. Based on the BIC criterion, there were three main clusters within the PAV matrix, the first being likely false positives (mean observance 2.8%; n=163,199). The second and third clusters were combined (mean observance 33%; n=33,350) and were used to calculate a distance matrix (method-jaccard) for visualizing subpopulation differences on a PCA plot (Figure 2e).

### Population structure

Population structure for both SNP and PAV data was estimated using fastStructure^41^ and Admixture^42^. SNP markers were randomly subsetted to 50k by LD pruning (parameters: --indep-pairwise 50 50 0.5) in plink 1.9^87^, while shell genes (as determined by DAPC clustering) were extracted from the pan-genome. In both analyses, a single sample with a maximum membership coefficient (qi) of <0.7 was considered admixed. Only non-admixed samples from the SNP analysis were used for further analysis. For SNP markers, multidimensional scaling (MDS), identity by state (IBS), and LD estimates (parameters: --r2 --ld-window-kb 500 --ld-window-r2 0) were performed in plink 1.9.

### Extraction of GBS markers from diversity panel assemblies

To integrate previously published GBS data^67^ with our new sequencing data, we extracted the relevant GBS markers from the assembled genomes. The paired end sequences were demultiplexed with the sabre package (https://github.com/najoshi/sabre). Samples were aligned to the reference using bwa-mem and SNP were further called using Varscan 2.3.9 (minimum depth of three and variant allele depth of two). SNP with more than 20% missing data in the GBS data were removed, and then those remaining were merged with the diversity panel (598 samples with 8.58M markers). A common set of 55,360 SNP was obtained.

### LD Decay

The extent of LD for the population was determined as described by^88^. For this, first we extracted one SNP every 100bp using plink (--bp-space 100), and selected a random set of 200,000 markers. LD (r^2^) was calculated using plink (--ld-window 500 --ld-window-kb 2000). The r^2^ value was averaged every 100bp of distance. A nonlinear model was fit for this data in R, and the extent was determined as when the LD (r^2^) nonlinear curve reaches 0.2. Average LD was 100 Kbp. This distance defined the window size for searching for candidate genes in the GWAS analyses.

### Search for *copia* elements

The *copia38* sequence (6.7 kb) was extracted from the genomic sequence of Seita.5G087200 using repbase (https://www.girinst.org/repbase/). Both the *copia38* sequence and *SvLes1* (Sevir.5G085400) sequence (both genomic sequence and CDS) were aligned to each of the pan-genome assemblies (n=598) using blat (-noHead -extendThroughN). From the blat results, each *copia38* alignment was checked whether it fell within the bounds of the *SvLes1* locus.

### Environmental correlations

Climate data were obtained from WorldClim^52^ using the Raster package in R for each of the 577 samples that have geographical coordinates. Correlations were calculated and visualized using the corrplot R package. To account for correlations between the 19 bioclimatic variables, we performed Principal component analysis (PCA) using prcomp in R. The top three principal components that contribute most of the variance were used independently as response variables in the association analysis. GEMMA^89^ was used to identify the association of bioclimatic variables and each of the SNP using only kinship in one model and using both kinship and population structure in another model. For population structure estimation, we first LD-pruned the markers in plink (indep-pairwise 50 50 0.5) and selected 50000 random markers. PCA was estimated in plink and the top three components were used as covariates in the mixed model to control for population structure. The best model was evaluated using Quantile-Quantile (Q-Q) plots of the observed vs. expected –log10(p) values which should follow a uniform distribution under the null hypothesis. SNP with p-values less than Bonferroni correction were considered significant. Genes within 100Kbp of a significant marker were also considered significant.

### Signatures of selection and local adaptation

We employed two statistics for scanning the genome-wide data for signs of positive natural selection: integrated haplotype score (iHS^90^), and Tajima’s D^91^. iHS is based on comparing the extended haplotype homozygosity (EHH) score of the ancestral and derived allele of each marker. This test detects loci where natural selection is driving one haplotype to high frequency, leaving recombination little time to break up the linkage group. This was calculated using hapbin^92^. To find genomic regions associated with natural selection, we estimated the fraction of SNP that have |iHS|>2.0 in each 100kbp window, with a slide of 10Kbp. The windows with the highest fraction are considered outliers. Tajima’s D^91^, compares the average number of pairwise differences (π) and the number of segregating sites (S). A negative value indicates positive selection. These values were calculated for 10k windows (with a slide of 10k) and the bottom 1% of the windows were considered outliers.

### GO and KEGG pathway enrichment analysis

GO enrichment analysis of positively selected genes was performed using topGO^93, 94^, an R Bioconductor package, to determine overrepresented GO categories across biological process (BP), cellular component (CC) and molecular function (MF) domains. Enrichment of GO terms was tested using Fisher’s exact test with p<0.05 considered as significant. KEGG^95^ pathway enrichment analysis was also performed on those gene sets based on a hypergeometric distribution test and pathways with p<0.05 were considered as enriched.

### GWAS and validation of *SvLes1*

The GWAS population to assess seed shattering was planted in the greenhouse facility at Donald Danforth Plant Science Center in April 2014. 215 accessions were chosen from the panel to perform the experiment (Table S8), with four replicates per accession. Shattering phenotype was measured by observing the amount of seed shattering after hand shaking of senesced dry plants. Individual plants were scored using a qualitative scale from 1 to 7. Genotypes were filtered at minor allele frequency > 5% for this population. GWAS was performed using a univariate mixed linear model from GEMMA^96^, with centered kinship matrix. We used the Wald test p-value^96^ for assessing significant peaks, but other p value estimates give similar results. SNP effects were identified using snpEff^82^. Deleterious effects of missense SNPs were predicted using PROVEA ^53^ on both the reference and alternative allele against the NCBI nr protein database.

To knockout *SvLes1* we used the backbone pTRANS_250d as described by Cermák et al.^97^. The protospacer of the guide RNAs targeted the first and second exons of *SvLes1*, upstream of the predicted causal mutation to ensure knock out by frameshift (Figure S12a). The binary vector was introduced into callus tissue using AGL1 agrobacterium. Tissue culture and transformation followed an established protocol for *S. viridis*^98^. T0 and T1 individuals were genotyped to identify newly acquired mutations near the targeted sites. A T2 homozygote *SvLes1-CRISPR1* was obtained and confirmed by Sanger sequencing, together with homozygotes of the unedited reference line for comparison. To test whether the non-shattering phenotype could be attributed to a single gene, we grew out the T3 seed from two presumptive T2 heterozygotes and several presumptive homozygotes and assessed their genotype at *SvLes1* by PCR and sequencing. Three inflorescences per plant were bagged at heading, and the bags left on until about half the seeds in the panicle appeared mature. Seeds that had fallen off in the bag were collected and weighed. Weights fell largely into two categories, either <35 mg, or >195 mg, with only a few weights in between, consistent with the effect of single gene.

### Germination rate, seed weight, and dispersal distance

Seeds from each genotype were collected, incubated at −80°C overnight, and then chlorine gas sterilized for 4 hours in a bell jar. Glumes were manually removed, and seeds were sowed onto 0.5X MS with 1% sucrose plates and incubated in the dark at 4°C for two days. Plates were moved into a growth chamber [12 hours light (156 µmol•m^-2^•sec^-1^) /12 hours dark, 31°C days/22 °C nights] and percent germination (judged by the emerging radicle tip) was recorded every 24 hours.

Seeds were harvested and pooled from five independent plants of each genotype. Five independent replicates of 20 seeds were weighed and recorded.

Starting at one week after heading and continuing to maturity, four random plants each from each of two sibling families of *SvLes1-CRISPR1* and wild-type *SvLes1-1* (ME034V) were individually put into a specially designed pot holder in a wind tunnel custom-built for the Kellogg lab with seed collection bins approximately every 10 cm. Blower speed was set to 5 mph for five minutes. Seeds collected in bins at various distances were counted to measure dispersal distance from the parent plant.

### Tensile strength measurement

Seeds of *SvLes1-1* (ME034v), *SvLes1-2* (A10.1), and *SvLes1-CRISPR1* were treated with 5% liquid smoke overnight at room temperature and kept in wet moss at 4°C in the dark for 2-3 weeks. Seeds were sown in Metro mix 360 and grown in a greenhouse with a 14h light/10 h dark cycle, day/night temperatures of 28 and 22°C and relative humidity of 40-50%. Panicles from main stems were collected at 8, 10, 12, 14, 16, 18, 20, 24 and 29 days after heading (the apex of the panicles emerged from the leaf sheath). Tensile strength of the spikelet and pedicel junction was measured as described previously^54^. Briefly, panicles were hung upside down from a Mark-10 model M3-2 force gauge. Spikelets were pulled off individually from a panicle using forceps and the peak tension was recorded. Only the most developed spikelets from the central third of the panicle were used to minimize the effects of developmental variation of the spikelets. Six plants with 20 spikelets from each plant were used per genotype per day of measurement. For *SvLes1-1* and *SvLes1-CRISPR1*, the plants in each genotype were offspring of two individual parent plants with the same allele.

### Histology

Histological procedures followed ^99^. Specifically, primary branches were collected from the central third of panicles 12 and 16 days after heading and fixed in FAA (37% formaldehyde: ethanol: H_2_O: acetic acid = 10:50:35:5), followed by a dehydration series in 50%, 70%, 85%, 95%, 100%, 100% and 100% ethanol and 25%, 50%, 75%, 100%, 100% and 100% Histo-Clear (National Diagnostics) series with ethanol as solvent. Paraplast (Leica Biosystems) was then added to each vial of samples and kept overnight, heated at 42°C, and placed in a 60°C oven. The solution was replaced with molten Paraplast twice a day for 3 days. Samples were then embedded in paraffin using a Leica EG1150 tissue embedder, sectioned in 10 µm serial slices with a Leica RM2255 automated microtome, and mounted on microscope slides at 42°C on a Premiere XH-2001 Slide Warmer. Sections were then deparaffinized, rehydrated, stained with 0.05% (w/v) toluidine blue O for 1.5 min, and then rinsed with water, dehydrated in ethanol, cleared with xylene and mounted with Permount Mounting Medium (Electron Microscopy Sciences)^36^. Images were taken using a Leica DM750 LED Biological microscope with ICC50 camera module and Leica Acquire v2.0 software.

### Domestication selective sweep

Raw sequencing reads of foxtail millet lines were obtained from a previous study^59^. Because the average sequencing coverage in the earlier study (∼0.5X) is much lower than in our study, we chose 79 lines (Table S16) that have an estimated coverage > 1X to maximize overlapping SNPs and perform analysis. Briefly, *S. italica* sequences were quality trimmed using sickle^100^, and aligned with bwa-mem to our *S. viridis* A10.1 genome. Multi-sample SNP calling was performed using samtools and Varscan with a minimum depth of 3. For *S.viridis*, the imputed, phased vcf was used for calculation of π, which uses high coverage. π calculation excluded missing samples. Shared SNPs between foxtail millet and *S. viridis* were combined, and missing data were imputed using Beagle 5.0^101^. Nucleotide diversity values π_viridis_ and π_italica_ were then calculated using vcftools^102^ at 100 kb window size. Using genome-wide nucleotide diversity as a reference, we used the program ms^103^ to conduct 100,000 coalescent simulations to estimate the variation range of π_italica_/π_viridis_ under a domestication bottleneck model for a window of 20 kb. Strength of the bottleneck was determined by genome wide π_italica_/π_viridis_. The estimated ranges were then compared to observed values π_italica_/π_viridis_ to determine significance of domestication selective sweep regions.

### Retrotransposon insertion in *S. italica Les1*

*Copia38* sequence was obtained from the foxtail millet genomic sequence^104^ near the ortholog of *SvLes1*, Seita.5G087200 (Si003873m.g). We confirmed the identity of *Copia38* and identified its LTR region by searching its sequence against repbase (https://www.girinst.org/repbase/). We used blastn^105^ to identify close homologs of *Copia38* in the Yugu1^104^ and A10.1 genomes. RaxML 8.2.9^106^ was used to construct the phylogeny of *Copia38* homologs, and pairwise distances of close homologs to *Copia38* were calculated using Kimura 2 parameter model. Read mapping to Yugu1 genome follows similar procedures described previously. Paired end reads spanning beyond the left and right junction point of *Copia38* were used to determine if the insertion occurs in an accession or not (Figure S12b).

### BSA mapping for small leaf angle

The cross between TB159 and A10.1 used pollen of TB159 and follows the protocol described in ^107^. F1 individuals were naturally self-pollinated to generate an F2 population. 417 F2 individuals were planted and phenotypically scored, and DNA from 30 small leaf angle individuals was pooled and sequenced. Sequences are available in the SRA at NCBI, BioProject number PRJNA527194 (to be released after publication). The analysis follows the methods described in a previous BSA study in *S. viridis*^40^. Identification of disruptive mutations and missense mutations with deleterious effects follows the same approach described in our GWAS study. Syntenic orthology between *SvLg2* and *liguleless2* in maize was examined and confirmed based on ^108^.

## Acknowledgements

We thank Zhonghui Wang, Xiaoping Li, and Hui Jiang for their help in maintaining the diversity panel and data collection, Dave Kudrna and Rod Wing at Arizona Genomics Institute for high molecular weight DNA extractions, and members of the Miller Lab at the Danforth Plant Science Center for helpful comments on the manuscript.

This work was supported by NSF grants DEB-0115397, MCB-0110809, DEB-0108501, PGRP-0952185, and IOS-1557633 to EAK; and DE-SC0008769 to TPB. The work conducted by the U.S. Department of Energy Joint Genome Institute is supported by the Office of Science of the U.S. Department of Energy under Contract No. DE-AC02-05CH11231.

## Author contributions

Credit taxonomy: Conceptualization: IB, TPB, MF, PH, EAK, JS; Methodology: PH, JJenkins, JS; Investigation: CC, KB, MF, AH, PH, JG, JJenkins, JJohnson,TK, JTL, SM, AS, HS, SS, TS, RT, JW, YY; Resources: MF, PH, EAK; Writing - Original Draft: KB, PH, JJ, SM, EAK; Writing - Review & Editing: IB, TPB, MF, PH, EAK, JTL, SM, JS; Visualization: AH, PH, JJ, EAK, JTL; Supervision: IB, TPB, EAK, JS; Project Administration: TPB, EAK, JS; Funding Acquisition: TPB, EAK, JS. Detailed contributions: JS: WGS assembly & sequencing project lead; JJ: map integration, chromosome assembly, analysis; JG: sequencing of BES, QC projects; HS, TK, TS: PacBio sequencing; SS: annotation; MF, PH, EAK: plant collecting; IB, MF, AH, PH, SM: SNP analyses; AH: PAV analyses; JTL: synteny analyses; SM: GWAS of environmental variables; TPB, PH: cloning of *SvLes1*, BSA; PH, EAK, DAN, RT, JW,: phenotyping of *SvLes1*; YY: histology and plant development.

## Data availability

Sequences are available in the SRA at NCBI, BioProject number PRJNA527194 (to be released after publication). Seed stocks for diversity lines are available by contacting co-authors I. Baxter or E. A. Kellogg.

---

**Fig. S1.** Cumulative citations for *Setaria viridis*, 1960-present, from Scopus, accessed Jan. 2, 2020.

**Fig. S2.** Population distribution and structure of North American accessions. Population assignment based on SNPs. **a** distribution of all accessions; **b** distribution of non-admixed samples only; **c** STRUCTURE plot including all accessions.

**Fig. S3.** Over-represented GO categories and KEGG enrichment pathways for each subpopulation.

**Fig. S4.** Correlations among the bioclim variables^52^ for the 577 accessions for which latitude-longitude information was available.

**Fig. S5.** GWAS results using each of the first three environmental PCs as response variable. PC1 found no significant loci, PC2 found one, and PC3 found several.

**Fig. S6.** Loci showing significant signatures of selection for each population. Left column, proportion of SNP with |iHS| >2 for each of the population. Right column, Tajima’s D represented in -ve scale. 100 Kbp windows (slide 10 Kbp). Horizontal lines represent 95th and 99th percentiles for iHS and 1st and 5th percentiles for Tajima’s D.

**Fig. S7. a** Comparison of seed germination rates of *SvLes1-1* and *SvLes1-CRISPR1* lines. Three plates of Setaria seed with n≥ 53 were measured for germination and averaged. Error is standard deviation. **b**, Comparison of seed weight of *SvLes1-1* and *SvLes1-CRISPR1* lines. Twenty seed weight was measured from 5 independent plants. *SvLes1-CRISPR1* seeds weighed significantly less (p<0.0001), but this did not affect germination.

**Fig. S8.** *SvLes1-2* (A10.1), *SvLes1-1*, and *SvLes1-CRISPR1* have similar anatomical structures in the abscission zone. Spikelets from main panicles 12 days after heading, stained with 0.05% Toluidine blue O. a,b *SvLes1-2* (A10.1). c,d *SvLes1-1.* e,f *SvLes1-CRISPR1.* a,c,e Scale bars = 100 µm. b,d,f Scale bars = 20 µm. En, endosperm; em, embryo. White dotted line, approximate position of AZ.

**Fig. S9.** Phylogenetic tree of *Copia38* copies in A10.1 and Yugu1 genomes. Red clade shows the recent explosion that contains the copy inserted into *SiLES1*. Gene models in green, *S. italica*; in blue, *S. viridis*.

**Fig. S10.** Representative dot plots**. a** Dot plot of clone 114809 on a region of scaffold_1. This alignment is representative of the high quality clone alignments in 239 of the 365 available clones. **b** Dot plot of clone 113526 on a region of scaffold_3, which is representative of the 36 clones that landed in repetitive regions of the genome.

**Fig. S11.** Plots of marker placements for each of the 9 chromosomes (scaffolds) of *S. viridis*.

**Fig. S12. a** guide RNA protospacers and their position relative to allelic mutations and gene model. **b** Method used to detect *Copia38* insertion.

**Table S1.** Metadata for 598 diversity lines. LIB, library; Q4 max, membership coefficient obtained from the output of STRUCTURE; Q4_subpop, subpopulation assignment using SNP data; Name, collector’s name and number or other unique identifier; new name, name in germplasm collection, lat, latitude of original collection; long, longitude of original collection. Subpopulation assignments correspond to those in figures, viz. Magenta -Central - 95 samples; Orange - West coast - 222 samples; Green – Central-North - 148 samples; blue - Central-East - 133 samples. bio1-bio19 are values of environmental variables from the WorldClim database ^52^. SRA is the accession number for the Sequence Read Archive at the NCBI.

**Table S2.** Subpopulation statistics. Numbers of non-admixed individuals based on SNP analysis; numbers used for each subpopulation in the PAV analysis are lower because not all libraries assembled well enough to be used. Number of over-represented genes is the number significant for that subpopulation alone. SNP, single nucleotide polymorphism; Pi/bp, polymorphisms per basepair; IBD, identity by descent for the subpopulation; IBS, identity by state for the subpopulation; LD, linkage disequilibrium; LD R^2^>=0.7%, percent of pairwise comparisons with LD>=0.7.

**Table S3** PAV diversity by subpopulation, listing over- and under-represented loci.

**Table S4.** Loadings of bioclim variables on the first three principal components. PC, principal component; min, minimum; max, maximum.

**Table S5.** Annotation of genes containing markers significantly associated with environmental variables.

**Table S6.** Significant Biological process GO terms for environmental associations.

**Table S7.** GO terms and KEGG pathways enriched for loci under selection, as identified by Tajima’s D and the absolute value of iHS.

**Table S8.** Setaria viridis samples scored for shattering phenotype. LIB, library number; Name, original accession identified; New_name, internal Brutnell lab identifier; Population, population assignment; Score_lowShatter, shattering index; Allele, nucleotide at position Chr_05:6849363.

**Table S9.** Comparison of pairwise diversity in the region surrounding *SiLes1* vs. *SvLes1*. Diversity is significantly lower in the *Les1* region in *S. italica* than in *S. viridis*, *p*=0.0066 based on 100,000 coalescent simulations. Description, interval considered for diversity estimates; total, total number of bp, centered on the gene; either side, number of bp upstream and downstream of the gene.

**Table S10.** Comparison of linkage disequilibrium (LD) in the region surrounding *SiLes1* vs. *SvLes1*. Intervals as in Table S9.

**Table S11.** Genomic libraries included in the *Setaria viridis* genome assembly and their respective assembled sequence coverage levels in the final release.

*Average read length of PACBIO reads.

**Table S12.** PACBIO library statistics for the libraries included in the *Setaria viridis* genome assembly and their respective assembled sequence coverage levels.

**Table S13.** Summary statistics of the initial output of the QUIVER polished MECAT assembly. The table shows total contigs and total assembled base pairs for each set of scaffolds greater than the size listed in the left hand column.

**Table S14.** Final summary assembly statistics for chromosome scale assembly. Final summary assembly statistics for chromosome scale assembly. Scaffold sequence total is all bases in the release plus gaps. Chromosome sequence is all bases in the chromosomes not including gaps. Total bases is all bases in the release excluding gaps.

**Table S15.** PAV matrix for all non-admixed individuals.

**Table S16.** *Setaria italica* lines used for π *italica* - π *viridis* comparison and for test of selective sweeps. Data from ^59^, downloaded from the Sequence Read Archive at NCBI.

