## Supplemental Figures for "The *Setaria viridis* genome and diversity panel enables discovery of a novel domestication gene"

### Slide 1
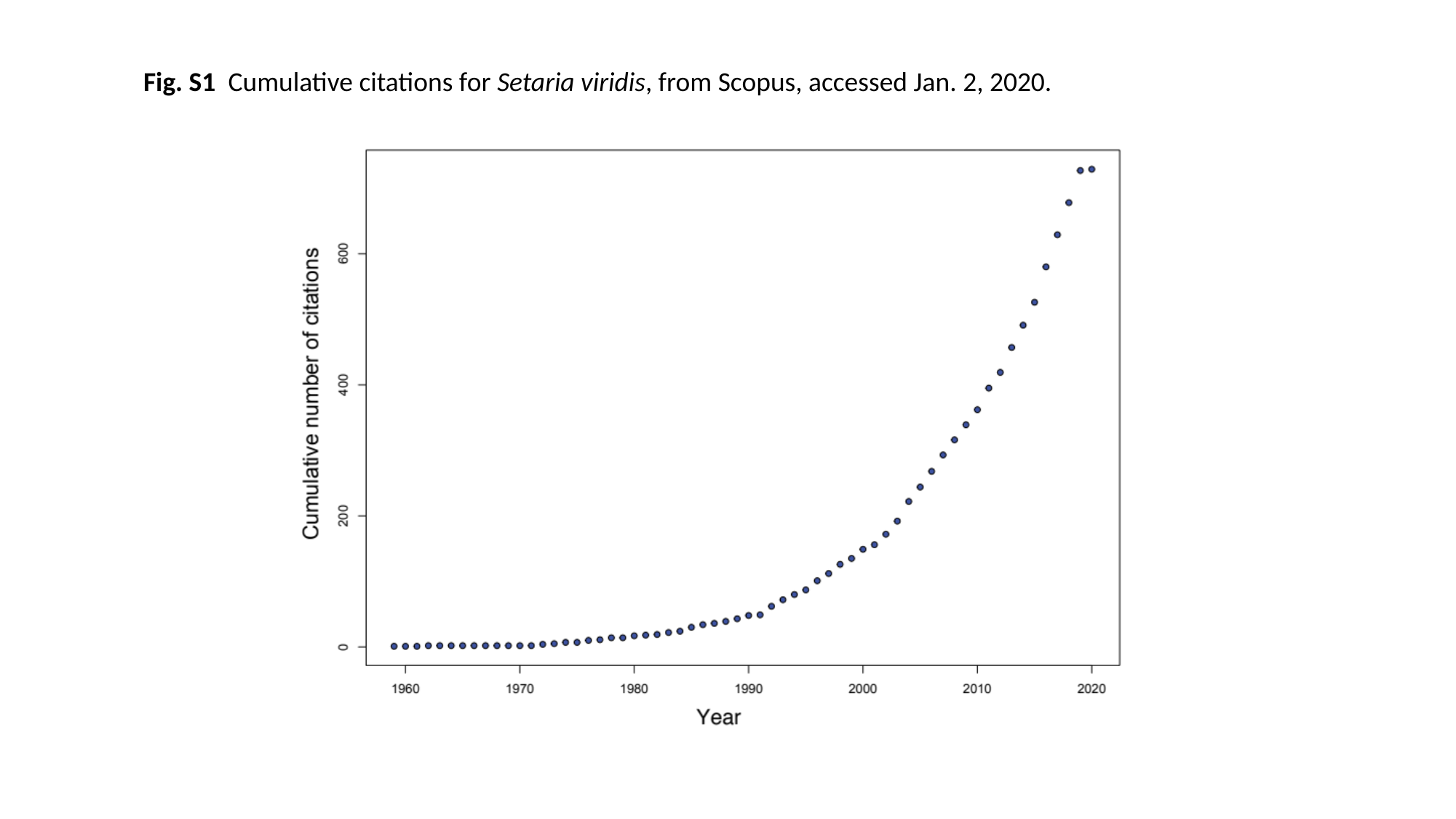

Fig. S1 Cumulative citations for Setaria viridis, from Scopus, accessed Jan. 2, 2020.

### Slide 2
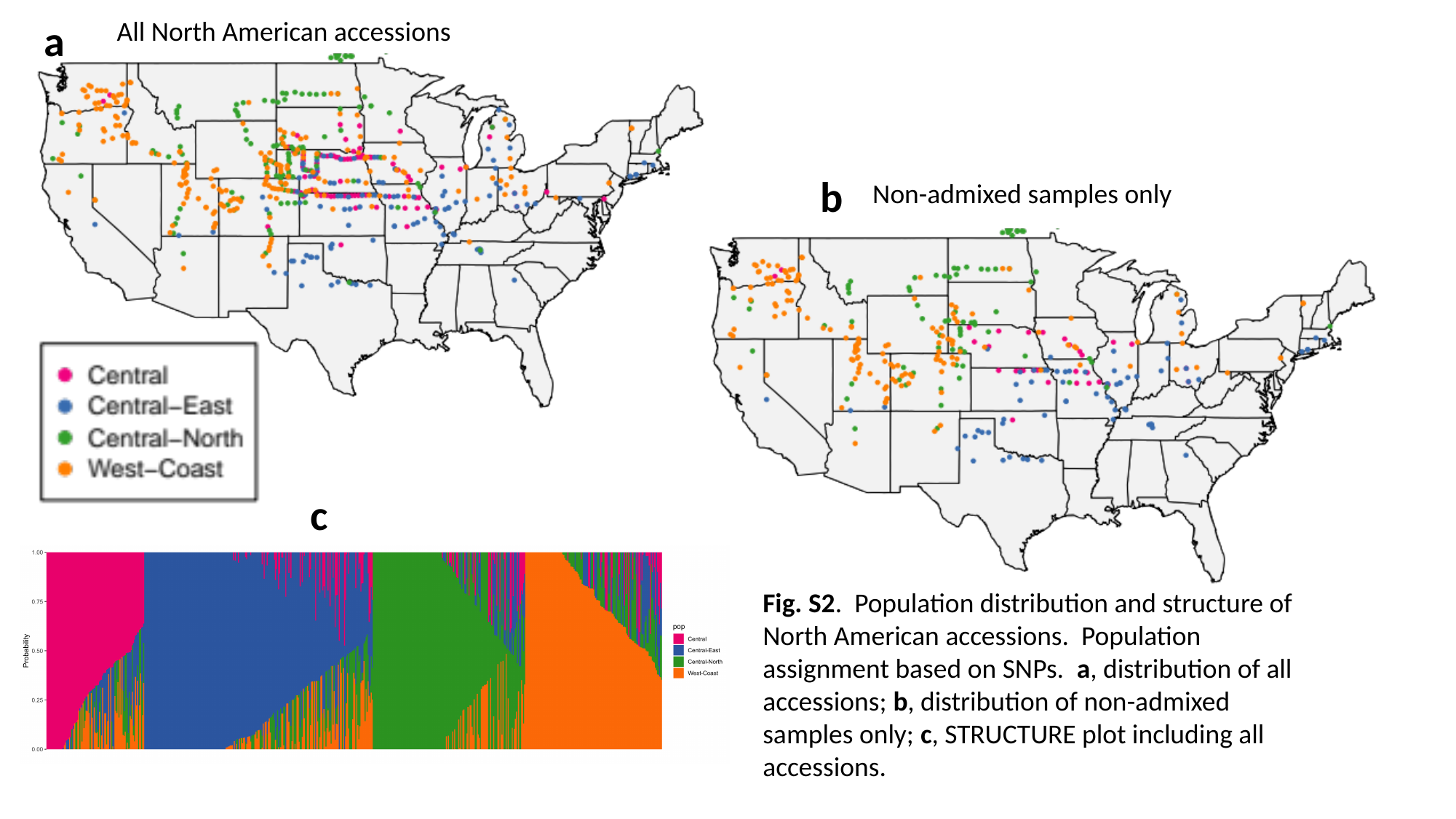

a
All North American accessions
b
Non-admixed samples only
c
Fig. S2. Population distribution and structure of North American accessions. Population assignment based on SNPs. a, distribution of all accessions; b, distribution of non-admixed samples only; c, STRUCTURE plot including all accessions.

### Slide 3
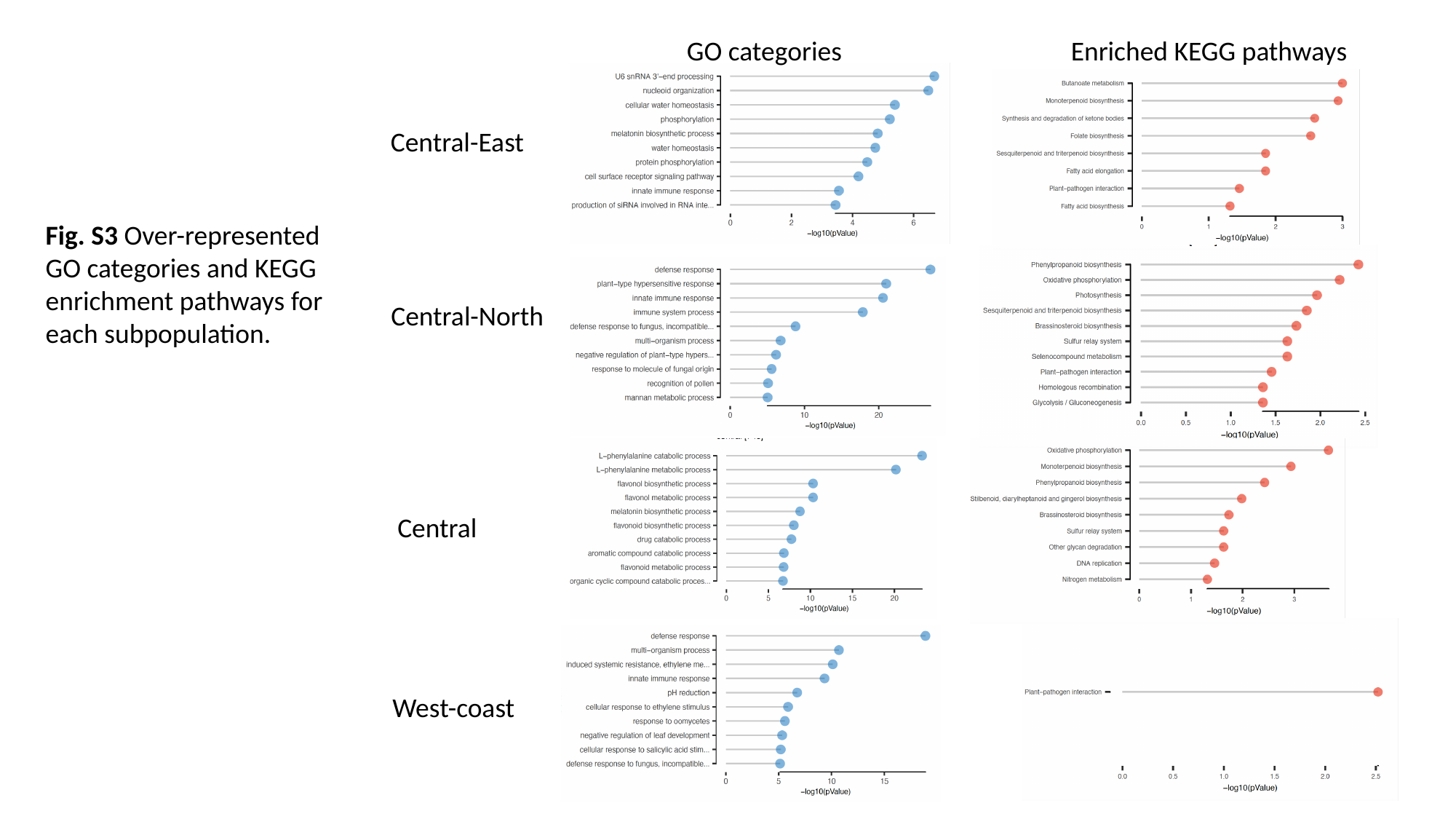

GO categories
Enriched KEGG pathways
Central-East
Fig. S3 Over-represented GO categories and KEGG enrichment pathways for each subpopulation.
Central-North
Central
West-coast

### Slide 4
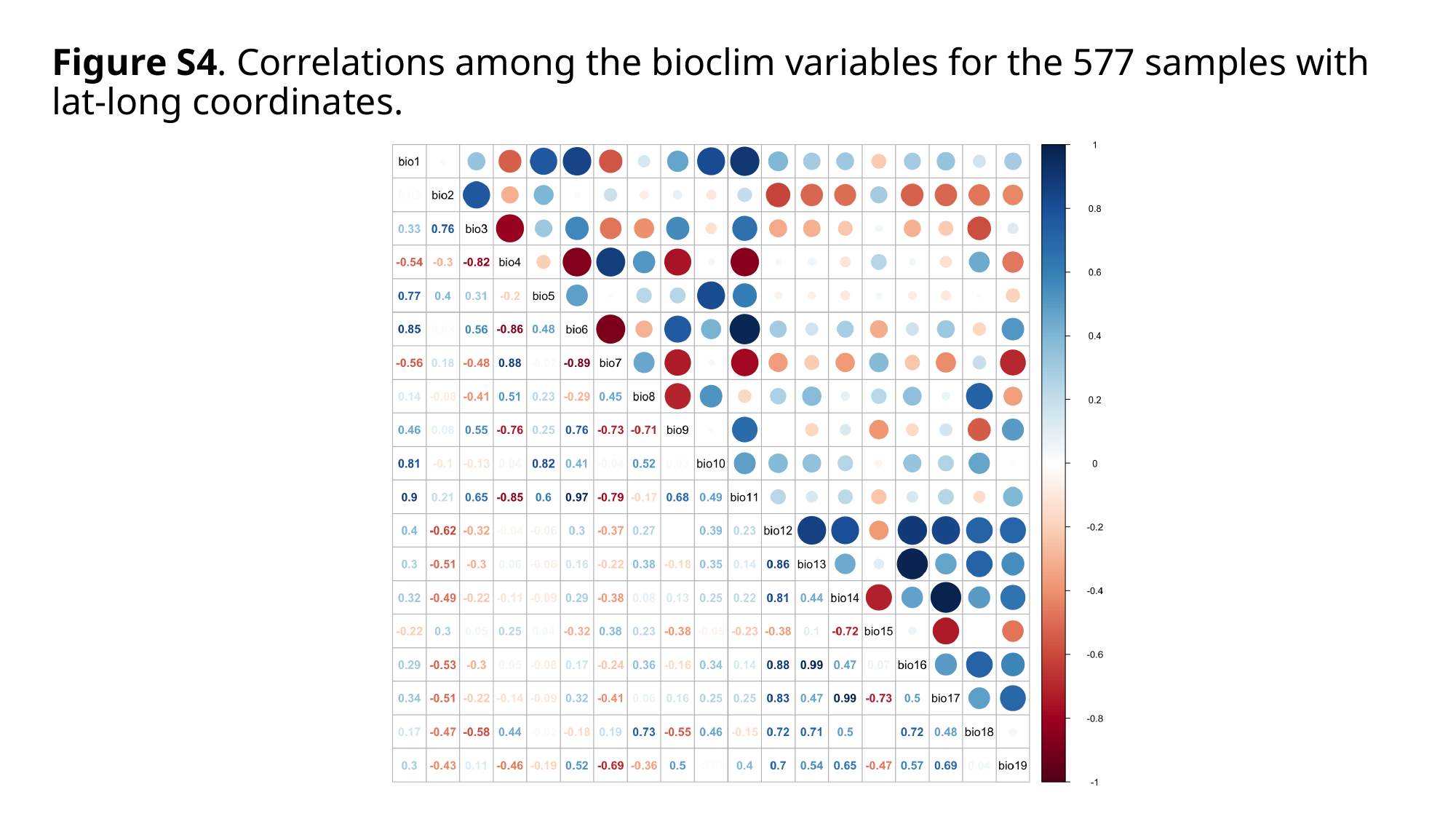

#### Figure S4. Correlations among the bioclim variables for the 577 samples with lat-long coordinates.

### Slide 5
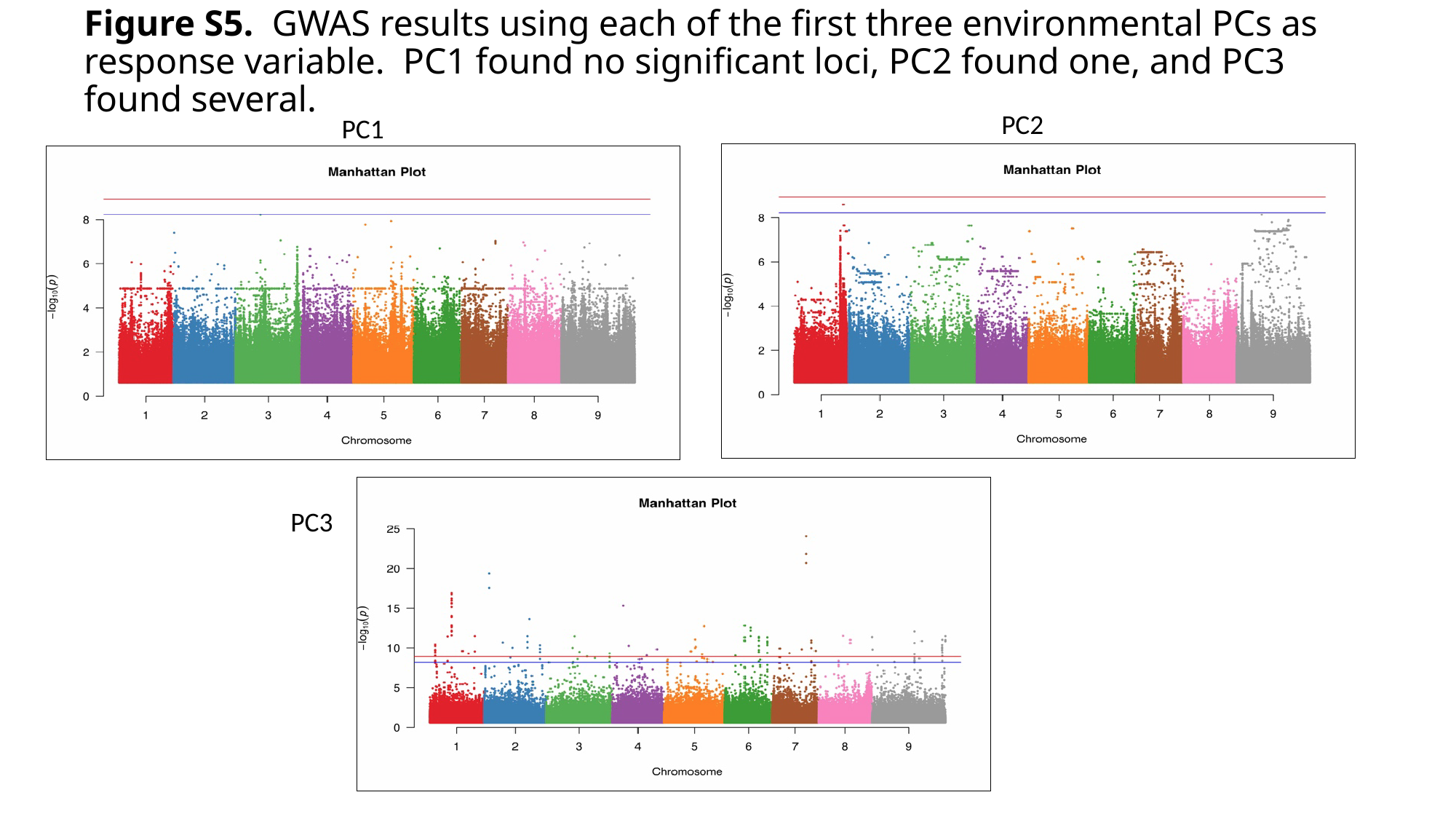

#### Figure S5. GWAS results using each of the first three environmental PCs as response variable. PC1 found no significant loci, PC2 found one, and PC3 found several.
PC2
PC1
PC3

### Slide 6
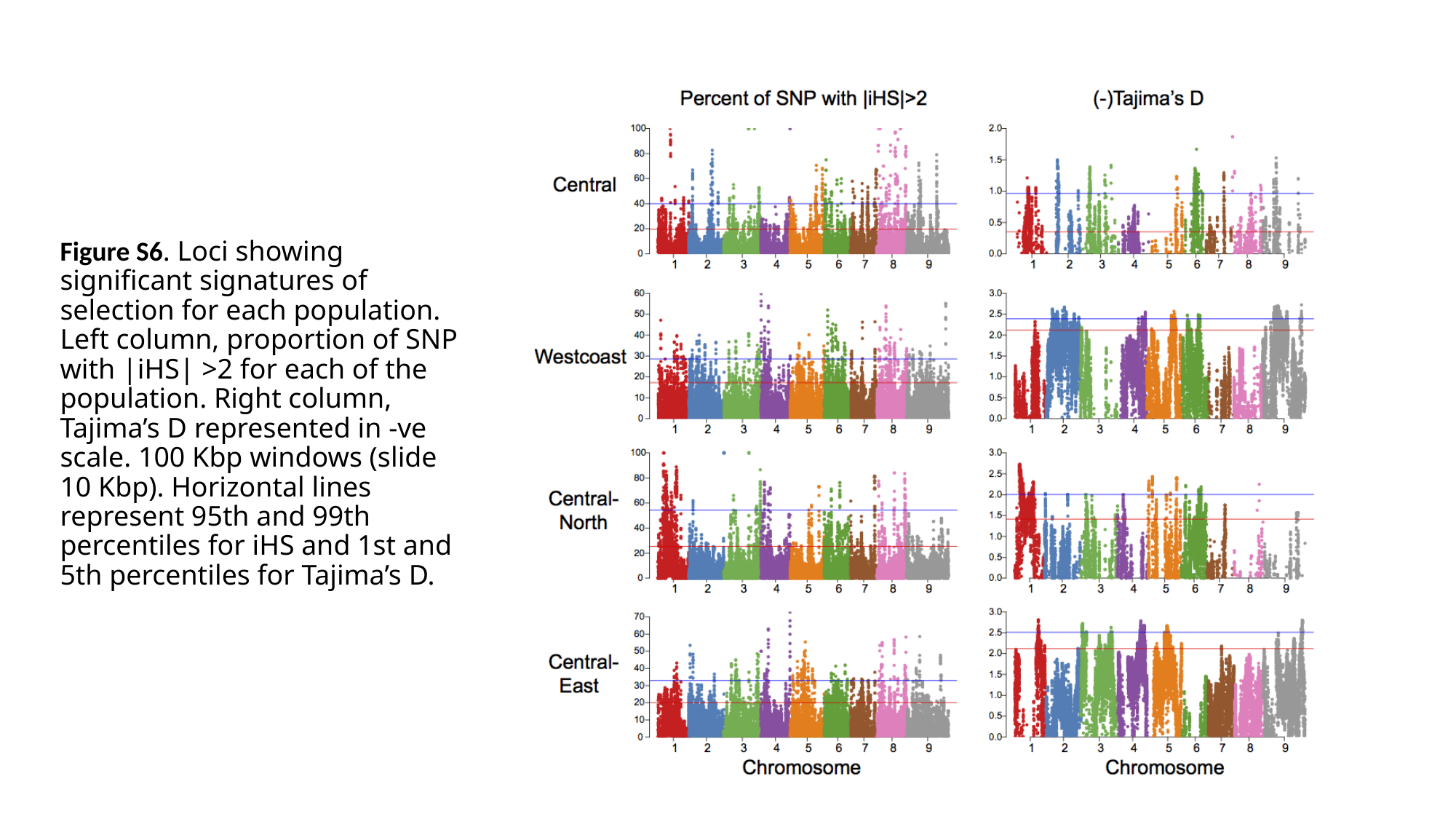

#### Figure S6. Loci showing significant signatures of selection for each population. Left column, proportion of SNP with |iHS| >2 for each of the population. Right column, Tajima’s D represented in -ve scale. 100 Kbp windows (slide 10 Kbp). Horizontal lines represent 95th and 99th percentiles for iHS and 1st and 5th percentiles for Tajima’s D.

### Slide 7
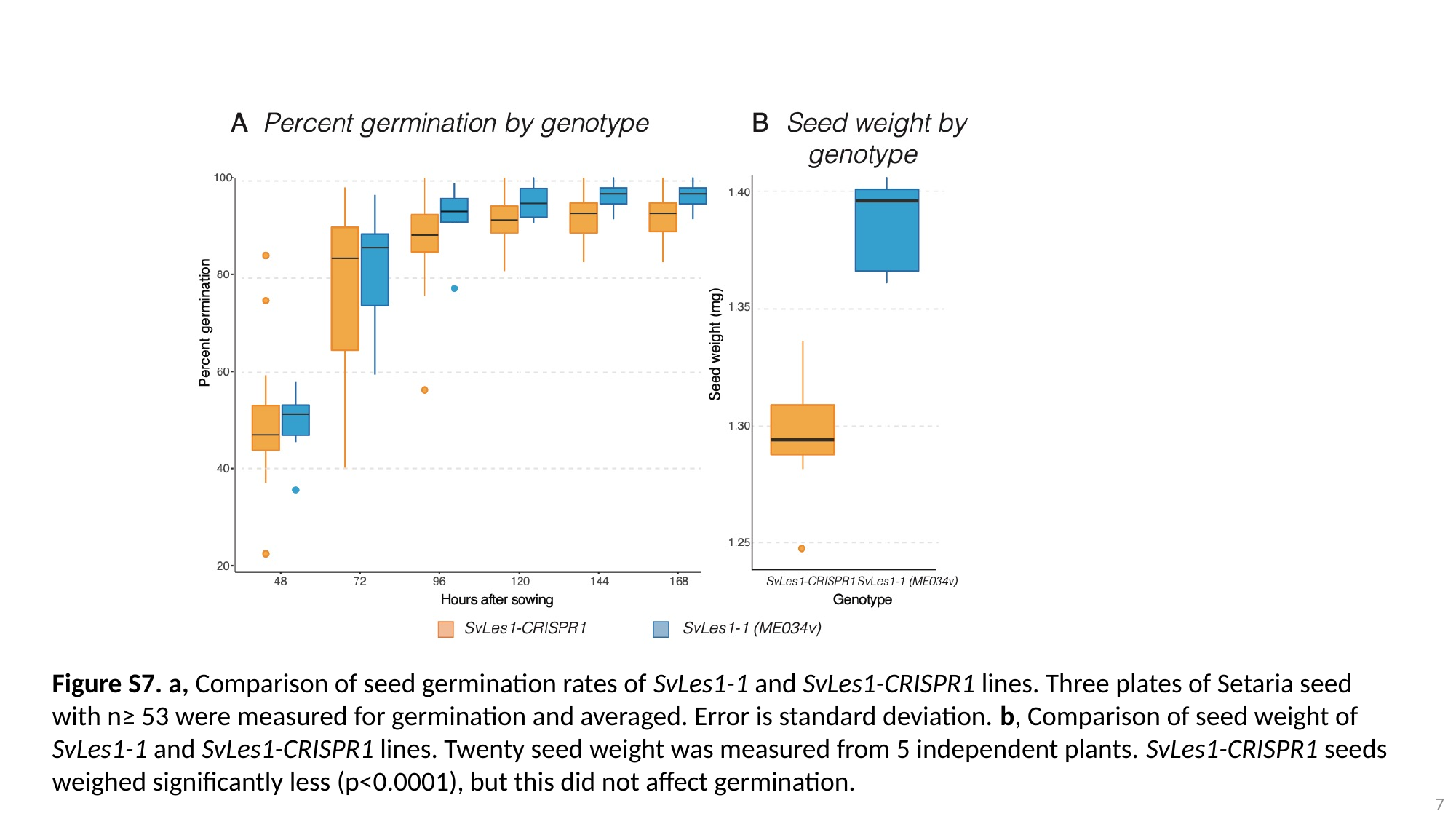

Figure S7. a, Comparison of seed germination rates of SvLes1-1 and SvLes1-CRISPR1 lines. Three plates of Setaria seed with n≥ 53 were measured for germination and averaged. Error is standard deviation. b, Comparison of seed weight of SvLes1-1 and SvLes1-CRISPR1 lines. Twenty seed weight was measured from 5 independent plants. SvLes1-CRISPR1 seeds weighed significantly less (p<0.0001), but this did not affect germination.
7

### Slide 8
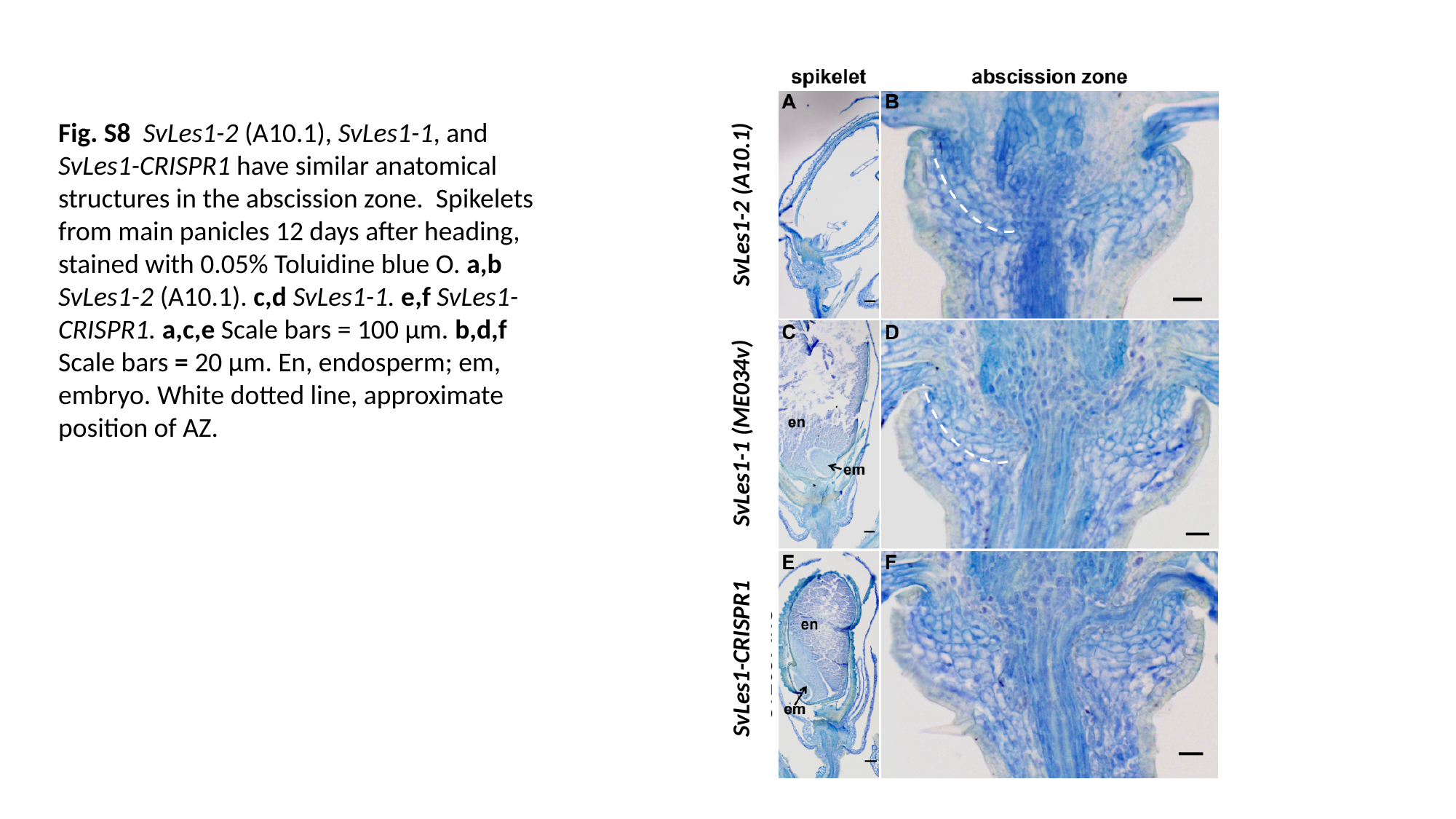

Fig. S8 SvLes1-2 (A10.1), SvLes1-1, and SvLes1-CRISPR1 have similar anatomical structures in the abscission zone. Spikelets from main panicles 12 days after heading, stained with 0.05% Toluidine blue O. a,b SvLes1-2 (A10.1). c,d SvLes1-1. e,f SvLes1-CRISPR1. a,c,e Scale bars = 100 µm. b,d,f Scale bars = 20 µm. En, endosperm; em, embryo. White dotted line, approximate position of AZ.
SvLes1-2 (A10.1)
SvLes1-1 (ME034v)
SvLes1-CRISPR1

### Slide 9
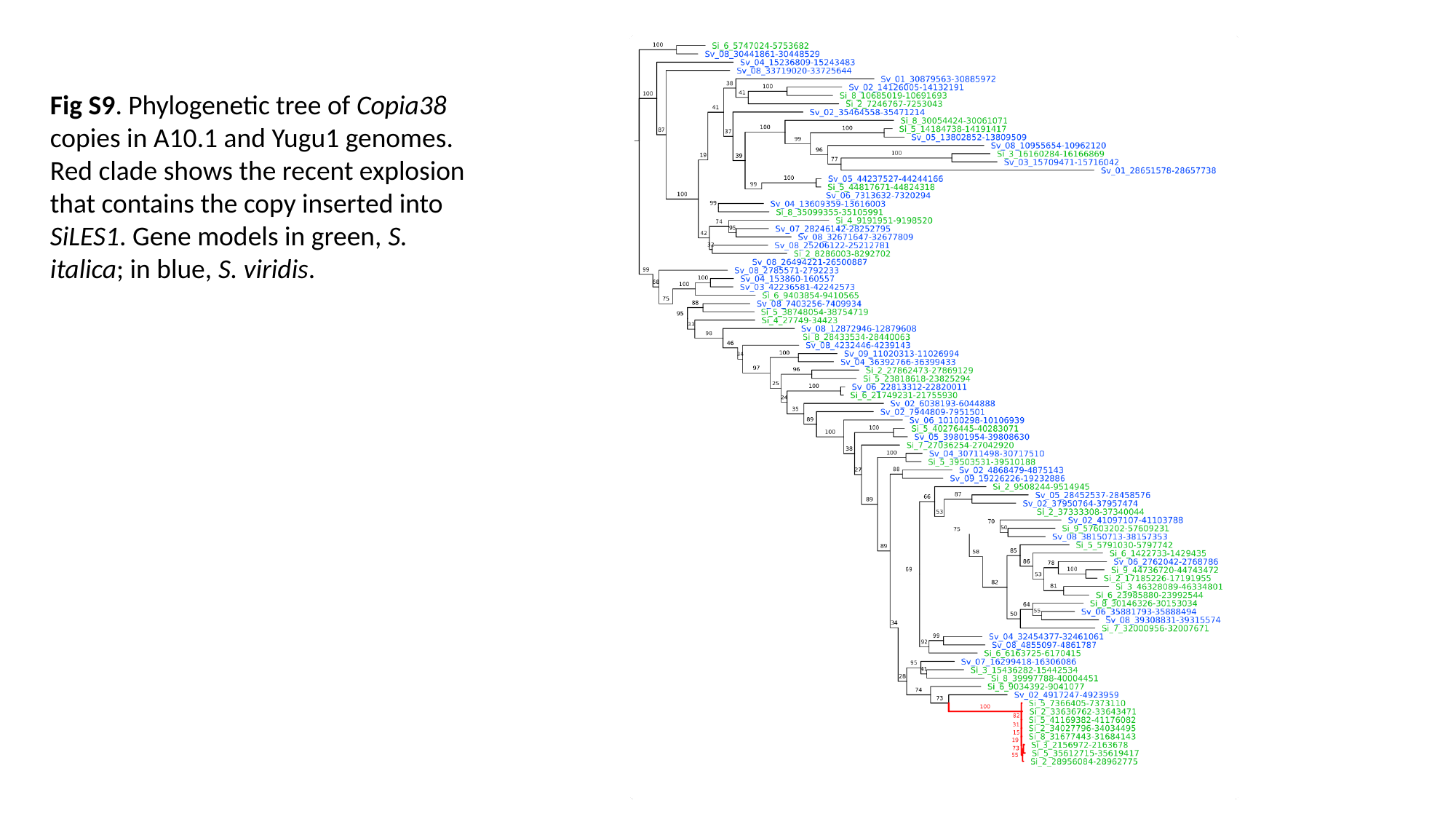

Fig S9. Phylogenetic tree of Copia38 copies in A10.1 and Yugu1 genomes. Red clade shows the recent explosion that contains the copy inserted into SiLES1. Gene models in green, S. italica; in blue, S. viridis.

### Slide 10
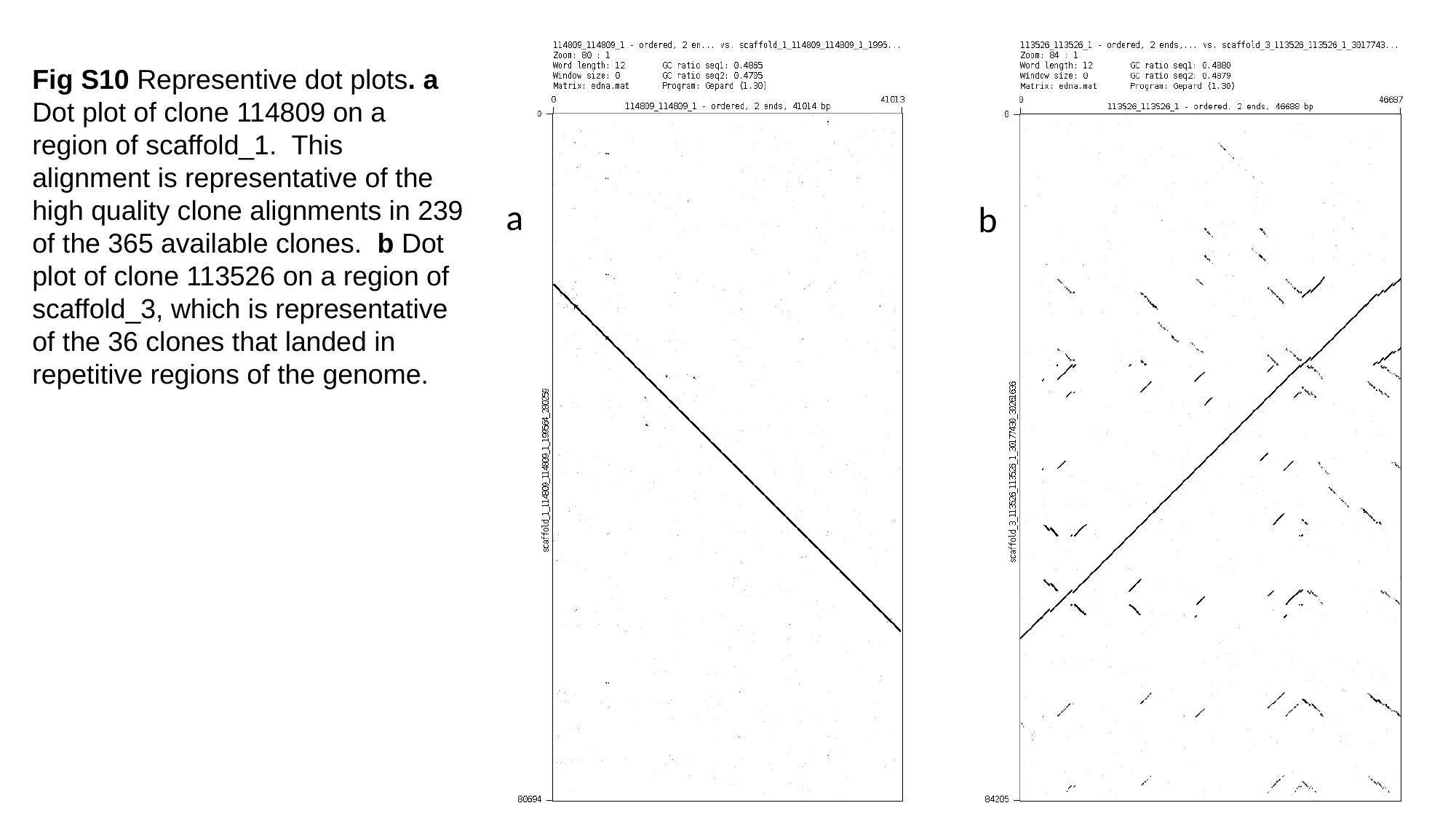

Fig S10 Representive dot plots. a Dot plot of clone 114809 on a region of scaffold_1. This alignment is representative of the high quality clone alignments in 239 of the 365 available clones. b Dot plot of clone 113526 on a region of scaffold_3, which is representative of the 36 clones that landed in repetitive regions of the genome.
a
b

### Slide 11
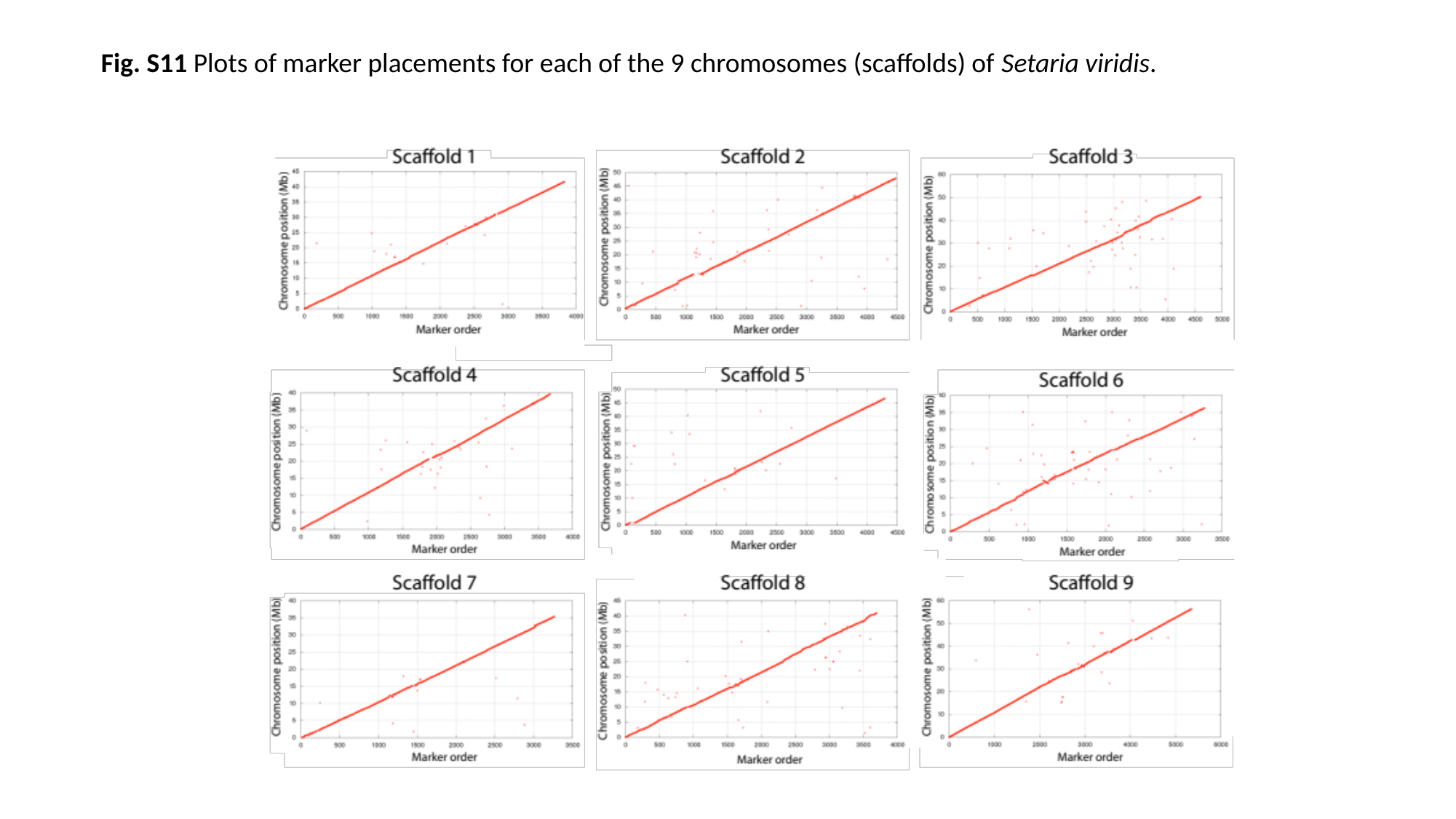

Fig. S11 Plots of marker placements for each of the 9 chromosomes (scaffolds) of Setaria viridis.

### Slide 12
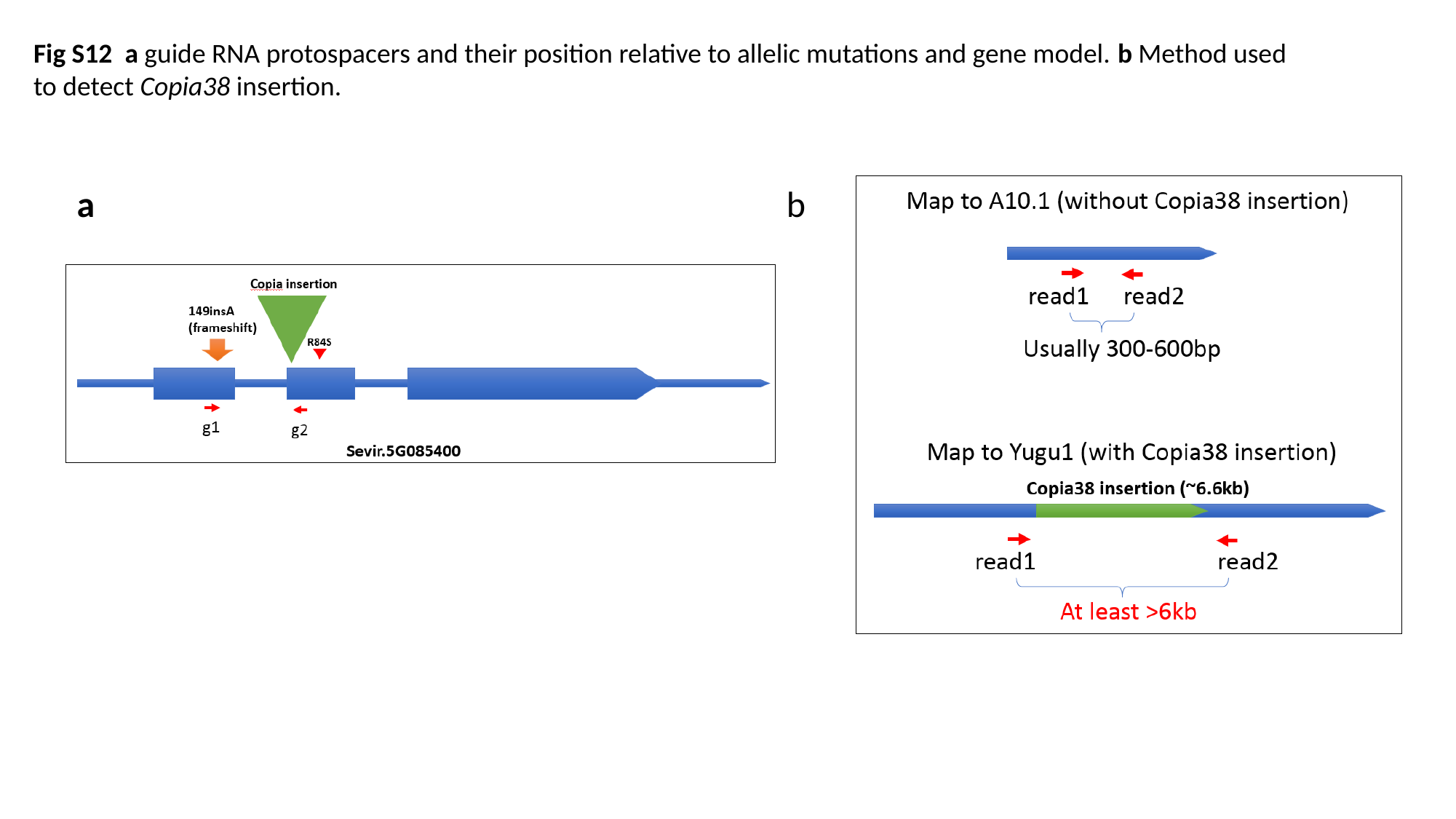

Fig S12 a guide RNA protospacers and their position relative to allelic mutations and gene model. b Method used to detect Copia38 insertion.
b
a
