## Supplemental Tables for "The *Setaria viridis* genome and diversity panel enables discovery of a novel domestication gene": Table S2.subpopulation.statistics.docx

Table S2. Subpopulation statistics. Numbers of non-admixed individuals based on SNP analysis; numbers used for each subpopulation in the PAV analysis are lower because not all libraries assembled well enough to be used. Number of over-represented genes is the number significant for that subpopulation alone. SNP, single nucleotide polymorphism; Pi/bp, polymorphisms per basepair; IBD, identity by descent for the subpopulation; IBS, identity by state for the subpopulation; LD, linkage disequilibrium; LD R^2^>=0.7%, percent of pairwise comparisons with LD>=0.7.

| Subpopulation | Central | Central East | Central North | Westcoast |
| --- | --- | --- | --- | --- |
| # non-admixed individuals | 44 | 81 | 104 | 153 |
| Unique SNPs | 836,585 | 21,747 | 368,608 | 183,712 |
| Pi/bp | 0.00457657 | 0.00235105 | 0.00373323 | 0.00259107 |
| #over-represented genes | 748 | 482 | 1904 | 435 |
| Subpopulation-specific private alleles | 24 | 0 | 20 | 1 |
| IBD - Average (kb) | 55.5021427 | 138.839981 | 162.239224 | 129.683235 |
| IBS_average | 8.2458E-05 | 1.6934E-05 | 2.3011E-05 | 1.998E-05 |
| LD mean | 0.2544468 | 0.4378058 | 0.3899508 | 0.4481402 |
| (LD R^2^>=0.7) % | 0.167 | 0.359 | 0.306 | 0.365 |
