## Supplemental Tables for "The *Setaria viridis* genome and diversity panel enables discovery of a novel domestication gene": Table S4.PC.loadings.docx

**Table S4**. Loadings of bioclim variables on the first three principal components. PC, principal component; min, minimum; max, maximum.

| Variable code | Variable name |  |  |  |
| --- | --- | --- | --- | --- |
|  |  | PC1 | PC2 | PC3 |
| bio6 | Min temperature warmest month | -0.326 | -0.191 | -0.089 |
| bio11 | Mean temperature coldest quarter | -0.305 | -0.204 | -0.185 |
| bio1 | Annual mean temperature | -0.300 | -0.072 | -0.324 |
| bio19 | Precipitation coldest quarter | -0.299 | 0.047 | 0.268 |
| bio12 | Annual precipitation | -0.278 | 0.276 | 0.037 |
| bio17 | Precipitation driest quarter | -0.277 | 0.191 | 0.162 |
| bio14 | Precipitation driest month | -0.264 | 0.194 | 0.155 |
| bio9 | Mean temperature driest quarter | -0.223 | -0.273 | 0.127 |
| bio16 | Precipitation wettest quarter | -0.203 | 0.281 | -0.049 |
| bio13 | Precipitation wettest month | -0.196 | 0.279 | -0.067 |
| bio10 | Mean temperature warmest quarter | -0.173 | 0.092 | -0.448 |
| bio5 | Max temperature warmest month | -0.108 | -0.127 | -0.468 |
| bio3 | Isothermality | -0.094 | -0.349 | -0.057 |
| bio18 | Precipitation warmest quarter | -0.078 | 0.364 | -0.168 |
| bio8 | Mean temperature wettest quarter | 0.048 | 0.260 | -0.361 |
| bio2 | Mean diurnal temperature range | 0.112 | -0.283 | -0.206 |
| bio15 | Precipitation seasonality | 0.196 | -0.010 | -0.240 |
| bio4 | Temperature seasonality | 0.245 | 0.288 | -0.058 |
| bio7 | Temperature annual range | 0.315 | 0.151 | -0.146 |
