## Supplemental Tables for "The *Setaria viridis* genome and diversity panel enables discovery of a novel domestication gene": Table S9. diversity estimates in italica vs. viridis.docx

**Table S9** Comparison of pairwise diversity in the region surrounding *SiLes1* vs. *SvLes1*. Diversity is significantly lower in the *Les1* region in *S. italica* than in *S. viridis*, *p*=0.0066 based on 100,000 coalescent simulations. Description, interval considered for diversity estimates; total, total number of bp, centered on the gene; either side, number of bp upstream and downstream of the gene.

| **Description** | **Start** | **End** | **Length (bp)** | **π *italica*** | **π *viridis*** | ***italica* SNP** | ***viridis* SNP** | **π *italica*/**  **π *viridis*** |
| --- | --- | --- | --- | --- | --- | --- | --- | --- |
| Gene | 6847970 | 6850236 | 2266 | 1.2650 | 4.4274 | 5 | 21 | 0.2857 |
| 10kb total | 6844103 | 6854103 | 10000 | 3.2352 | 50.1570 | 96 | 189 | 0.0645 |
| 20kb total | 6839103 | 6859103 | 20000 | 6.3833 | 135.5472 | 245 | 460 | 0.0471 |
| 50kb total | 6824103 | 6874103 | 50000 | 20.9668 | 286.5908 | 717 | 1001 | 0.0732 |
| 10kb either side | 6837970 | 6860236 | 22266 | 7.0845 | 159.3458 | 300 | 545 | 0.0445 |
| 20kb either side | 6827970 | 6870236 | 42266 | 20.3942 | 247.2280 | 640 | 871 | 0.0825 |
| 50kb either side | 6797970 | 6900236 | 102266 | 39.1945 | 959.0633 | 1129 | 3233 | 0.0408 |
