## Supplemental Tables for "The *Setaria viridis* genome and diversity panel enables discovery of a novel domestication gene": Table S10. Comparison of linkage disequilibrium.docx

**Table S10** Comparison of linkage disequilibrium (LD) in the region surrounding *SiLes1* vs. *SvLes1*. Intervals as in Table S9.

|  | | | | ***Setaria viridis*** | | | ***Setaria italica*** | | |
| --- | --- | --- | --- | --- | --- | --- | --- | --- | --- |
| **Description** | **Start** | **End** | **Length (bp)** | **Mean** | **% of comp. (>=0.5)** | **% of comp. (>=0.7** | **Mean** | **% of comp. (>=0.5)** | **% of comp. (>=0.7** |
| Gene | 6847970 | 6850236 | 2266 | 0.23 | 22.07 | 20.69 | 0.80 | 100.00 | 100.00 |
| 10kb total | 6844103 | 6854103 | 10000 | 0.23 | 19.37 | 14.60 | 0.96 | 100.00 | 100.00 |
| 20kb total | 6839103 | 6859103 | 20000 | 0.26 | 24.78 | 15.28 | 0.97 | 100.00 | 100.00 |
| 50kb total | 6824103 | 6874103 | 50000 | 0.29 | 27.17 | 20.68 | 0.30 | 25.91 | 25.91 |
| 10kb either side | 6837970 | 6860236 | 22266 | 0.27 | 25.63 | 17.36 | 0.98 | 100.00 | 100.00 |
| 20kb either side | 6827970 | 6870236 | 42266 | 0.28 | 25.63 | 19.51 | 0.30 | 25.91 | 25.91 |
| 50kb either side | 6797970 | 6900236 | 102266 | 0.27 | 22.68 | 16.64 | 0.46 | 42.35 | 42.13 |
