## Supplemental Tables for "The *Setaria viridis* genome and diversity panel enables discovery of a novel domestication gene": Table S11-S14.assembly.statistics.docx

**Table S11** Genomic libraries included in the *Setaria viridis* genome assembly and their respective assembled sequence coverage levels in the final release.

^*^Average read length of PACBIO reads.

| **Library** | **Sequencing Platform** | **Assembly**  **Release** | **Average Read/Insert Size** | **Read Number** | **Assembled Sequence Coverage (X)** |
| --- | --- | --- | --- | --- | --- |
| H0022 | Illumina | 2.0 | 800 | 425,635,116 | 240 |
|  | PACBIO | 2.0 | 10,587^*^ | 4,768, 857 | 118.18 |

**Table S12** PACBIO library statistics for the libraries included in the *Setaria viridis* genome assembly and their respective assembled sequence coverage levels.

| **Cutoff** | **Number of Reads** | **Basepairs** | **Average Read Length** | **Coverage** |
| --- | --- | --- | --- | --- |
| 0 | 4,768,857 | 59,091,859,660 | 10,587 | 118.18x |
| 1,000 | 4,427,654 | 58,922,852,295 | 11,597 | 117.85x |
| 2,000 | 4,128,899 | 58,477,661,257 | 12,522 | 116.96x |
| 3,000 | 3,875,244 | 57,847,410,625 | 13,346 | 115.69x |
| 4,000 | 3,653,717 | 57,073,702,495 | 14,086 | 114.15x |
| 5,000 | 3,442,812 | 56,125,591,515 | 14,820 | 112.25x |
| 6,000 | 3,238,591 | 55,003,123,843 | 15,551 | 110.01x |
| 7,000 | 3,041,062 | 53,719,962,620 | 16,270 | 107.44x |
| 8,000 | 2,850,014 | 52,287,501,523 | 16,979 | 104.58x |
| 9,000 | 2,665,092 | 50,716,192,061 | 17,681 | 101.43x |
| 10,000 | 2,486,248 | 49,017,650,265 | 18,373 | 98.04x |
| 11,000 | 2,314,198 | 47,211,798,377 | 19,049 | 94.42x |
| 12,000 | 2,148,024 | 45,301,509,765 | 19,727 | 90.60x |
| 13,000 | 1,990,291 | 43,330,677,398 | 20,385 | 86.66x |
| 14,000 | 1,839,724 | 41,298,472,587 | 21,043 | 82.60x |
| 15,000 | 1,696,209 | 39,218,281,557 | 21,692 | 78.44x |
| 16,000 | 1,557,328 | 37,065,930,006 | 22,354 | 74.13x |
| 17,000 | 1,422,360 | 34,839,361,404 | 23,034 | 69.68x |
| 18,000 | 1,291,347 | 32,547,099,163 | 23,740 | 65.09x |
| 19,000 | 1,163,251 | 30,177,734,218 | 24,475 | 60.36x |

**Table S13** Summary statistics of the initial output of the QUIVER polished MECAT assembly. The table shows total contigs and total assembled basepairs for each set of scaffolds greater than the size listed in the left hand column.

| **Minimum**  **Scaffold**  **Length** | **Number of**  **Scaffolds** | **Number of**  **Contigs** | **Scaffold Size** | **Basepairs** | **% Non-gap Basepairs** |
| --- | --- | --- | --- | --- | --- |
| 5 Mb | 25 | 25 | 347,994,520 | 347,994,520 | 100.00% |
| 2.5 Mb | 33 | 33 | 378,539,963 | 378,539,963 | 100.00% |
| 1 Mb | 40 | 40 | 389,332,751 | 389,332,751 | 100.00% |
| 500 Kb | 44 | 44 | 392,513,339 | 392,513,339 | 100.00% |
| 250 Kb | 49 | 49 | 394,501,069 | 394,501,069 | 100.00% |
| 100 Kb | 53 | 53 | 395,090,738 | 395,090,738 | 100.00% |
| 50 Kb | 77 | 77 | 396,626,132 | 396,626,132 | 100.00% |
| 25 Kb | 107 | 107 | 397,813,004 | 397,813,004 | 100.00% |
| 10 Kb | 110 | 110 | 397,864,082 | 397,864,082 | 100.00% |
| 5 Kb | 110 | 110 | 397,864,082 | 397,864,082 | 100.00% |
| 2.5 Kb | 110 | 110 | 397,864,082 | 397,864,082 | 100.00% |
| 1 Kb | 110 | 110 | 397,864,082 | 397,864,082 | 100.00% |
| 0 bp | 110 | 110 | 397,864,082 | 397,864,082 | 100.00% |

**Table S14** Final summary assembly statistics for chromosome scale assembly. Scaffold sequence total is all bases in the release plus gaps. Chromosome sequence is all bases in the chromosomes not including gaps. Total bases is all bases in the release excluding gaps.

| **Scaffold total** | 14 |
| --- | --- |
| **Contig total** | 75 |
| **Scaffold sequence total** | 395.7 Mb |
| **Chromosome Sequence** | 394.9 Mb |
| **Total bases** | 395.1 Mb |
| **Scaffold N/L50** | 4 / 46.7 Mb |
| **Contig N/L50** | 11 / 11.2 Mb |
